# Dynamic and causal contribution of parietal circuits to perceptual decisions during category learning

**DOI:** 10.1101/304071

**Authors:** Lin Zhong, Yuan Zhang, Chunyu A. Duan, Jingwei Pan, Ning-long Xu

**Affiliations:** Institute of Neuroscience, State Key Laboratory of Neuroscience, CAS Center for Excellence in Brain Science and Intelligence Technology, Shanghai Institutes for Biological Sciences, Chinese Academy of Sciences, Shanghai 200031, China.; University of Chinese Academy of Sciences, Shanghai 200031, China.

## Abstract

Making perceptual decisions to categorize unknown sensory stimuli is a fundamental cognitive function, known as category learning. The posterior parietal cortex (PPC), although has been intensively studied for its role in decision-making and other cognitive functions, its causal link with behavior remains controversial. Here we combine *in vivo* two-photon imaging, circuit manipulation and auditory psychophysics behavior in mice to probe the role of PPC and its connectivity with sensory cortex in decision-making during category learning. We show that PPC neuronal populations exhibit coding dynamics characteristic of category learning, showing representations for both new sensory stimuli and prior learned categories. Circuit-specific perturbations of PPC and its projections to auditory cortex impaired decision performance specifically for categorizing new auditory stimuli. These data reveal a dynamic and causal role of the parietal-auditory circuit in decision-making, integrating prior knowledge to guide categorical decisions on new sensory stimuli.

## Introduction

The ability of humans and other animals to classify unknown items into behaviorally relevant categories or concepts, known as category learning, is essential for survival and fundamental to higher cognition (*1–5*). For example, to decide a newly encountered animal as a predator or prey, or an unknown plant as nutrients or poisons, the subject needs to compare the perceptual properties of the unfamiliar items with prior experiences. Central to this process is making perceptual judgements on novel sensory stimuli based on their perceptual relationship to prior learned prototypes or exemplars, an essential step in category learning. The underlying causal neural circuit causal mechanism remains poorly understood. The posterior parietal cortex (PPC) has been widely reported to show decision and categorization-related activity both in primates (*6–12*) and in rodents (*13–17*), and has also been implicated in many other cognitive functions (*8,14,18–22*). Therefore, PPC and its connectivity with related regions are likely to be the key brain system in category learning. However, recent studies reported rather controversial results regarding the causal role of PPC in choice behavior. In macaques, inactivation of PPC did not have significant effect on behavioral performance in the random-dot visual motion decision task (*23*) or a self-motion perceptual decision task (*24*). In rodents, silencing PPC neither had significant behavioral effect in auditory or tactile decision tasks (*17, 25, 26*). But silencing or disruption of PPC activity impaired visually guided choice behavior in rodents (*14, 17, 27, 28*). Thus, it has become controversial whether PPC is functionally significant for general decisionmaking processes, or it is only required specifically in visual-related functions in rodents.

Most of these studies, however, focused on the PPC activity after animals were well-trained by the same set of sensory stimuli also being used for assessing performance following perturbation, whereas in the real world, animals make decisions upon unfamiliar sensory stimuli based on their relationship to prior learned exemplars or prototypes, a learning capability to survive dynamic and changing environments (*1–5*). We thus sought to determine the role of PPC in categorical decision-making upon new sensory stimuli that were not used for prior training. Combining mouse auditory psychophysics behavior with *in vivo* two-photon calcium imaging and circuit manipulations, we demonstrate that PPC and its connectivity with the auditory cortex are dynamically and causally involved in categorical decision-making on new sensory stimuli, providing a causal circuit mechanism for category learning.

## Auditory category learning in head-fixed mice

We trained head-fixed mice to perform a two-alternative-forced-choice (2AFC) auditory discrimination task using two exemplar tones (training stimuli, 2 octaves apart, 8 kHz and 32 kHz; **Fig. 1A**). Mice report their perceptual decisions on tone stimuli as “low” or “high” frequency by licking the left or right lick-port (**Fig. 1A**; see **fig. S1** and **movie S1**). We then introduced new “test” tones in randomly interleaved trials (30%) to examine whether mice can make categorical judgements on new sensory stimuli based on prior trainings (**Fig. 1A**). Remarkably, mice exhibit sharp categorical decisions for the test stimuli (**Fig. 1, B and C**), indicated by the peak discriminability (the slope value at the categorization boundary) significantly greater than that for an otherwise linear relation between choice and tone frequency (**Fig. 1, C and D**). Mice’s subjective categorization boundaries (points of subjective equality, PSE) are distributed in close proximity of experimentally defined category boundary (16 kHz; **Fig. 1E**; also see **fig. S1E**). These data indicate that mice are able to make categorical decisions to classify new tone stimuli as “high” or “low” based on previous trainings, a key process in category learning (*29*).

**Fig. 1.**
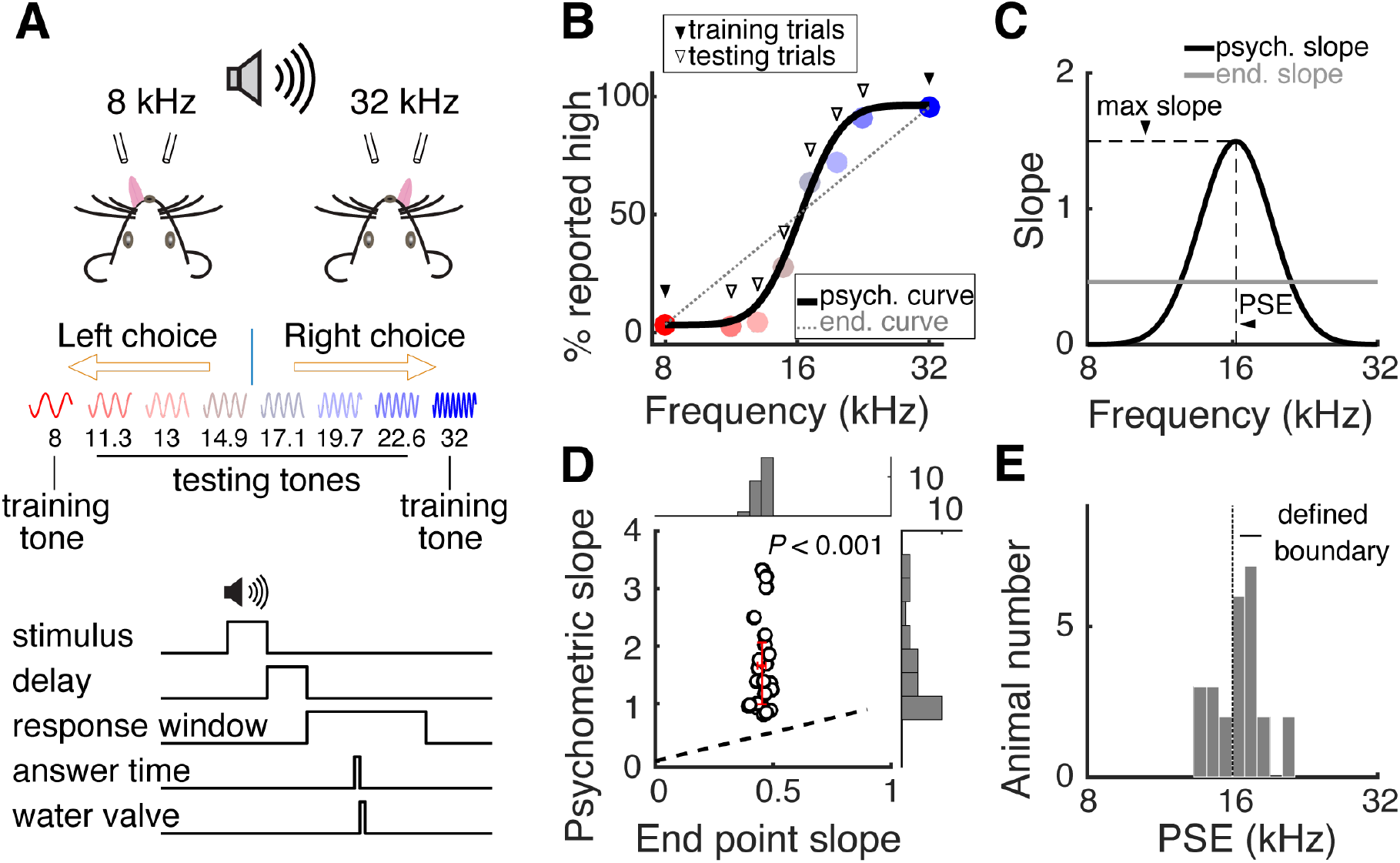
Perceptual categorization of auditory stimulus in head-fixed mice. **(A)** Schematic showing the configuration of behavioral paradigm and trial structure. **(B)** Psychometric function from an example behavioral session where the test stimuli were first introduced. **(C)** Slopes of the psychometric function and hypothetical linear relation between stimulus and behavioral choice shown in **(B)** The peak of the slope of the psychometric function (black) around the point of subjective equality (PSE) indicates perceptual categorization. The slope of the linear function (grey) shows a constant value. **(D)** Scatter plot of group data comparing peak psychometric slopes (1.66 ± 0.16) and linear relation slopes (0.45 ± 0.01) within each animal (P < 0.0001, n = 25 mice). **(E)** Histogram showing distribution of the PSE of 25 animals from the first session introducing test stimuli (PSE = 16.5 ± 0.3 kHz). Red error bars indicate median and interquartile range.

## PPC neurons exhibit characteristic dynamics of category learning

Although the dynamics of PPC activity has been recently studied (*13, 14, 16, 21, 30*), the precise relationship between PPC activity and specific learning process is still unclear. Here we investigate the relationship between PPC activity dynamics and perceptual category learning, using chronic *in vivo* two-photon calcium imaging from PPC neurons across different sessions/ days. Genetically encoded calcium indicator GCaMP6s (*31*) was expressed in PPC of left hemisphere using AAV vectors to indicate the intracellular calcium dynamics, followed by implantation of a chronic imaging window (*32*). The same population of layer 2/3 neurons were imaged during task performance repeatedly across days starting from the session when the test stimuli were first introduced (**Fig. 2A; fig. 2**; also see methods in supplementary materials). Consistent with previous studies (*14, 16, 22, 27*), we found that PPC neurons show complex response selectivity to multiple behavioral variables (**Fig. 2B; fig. 3; table S1**).

**Fig. 2.**
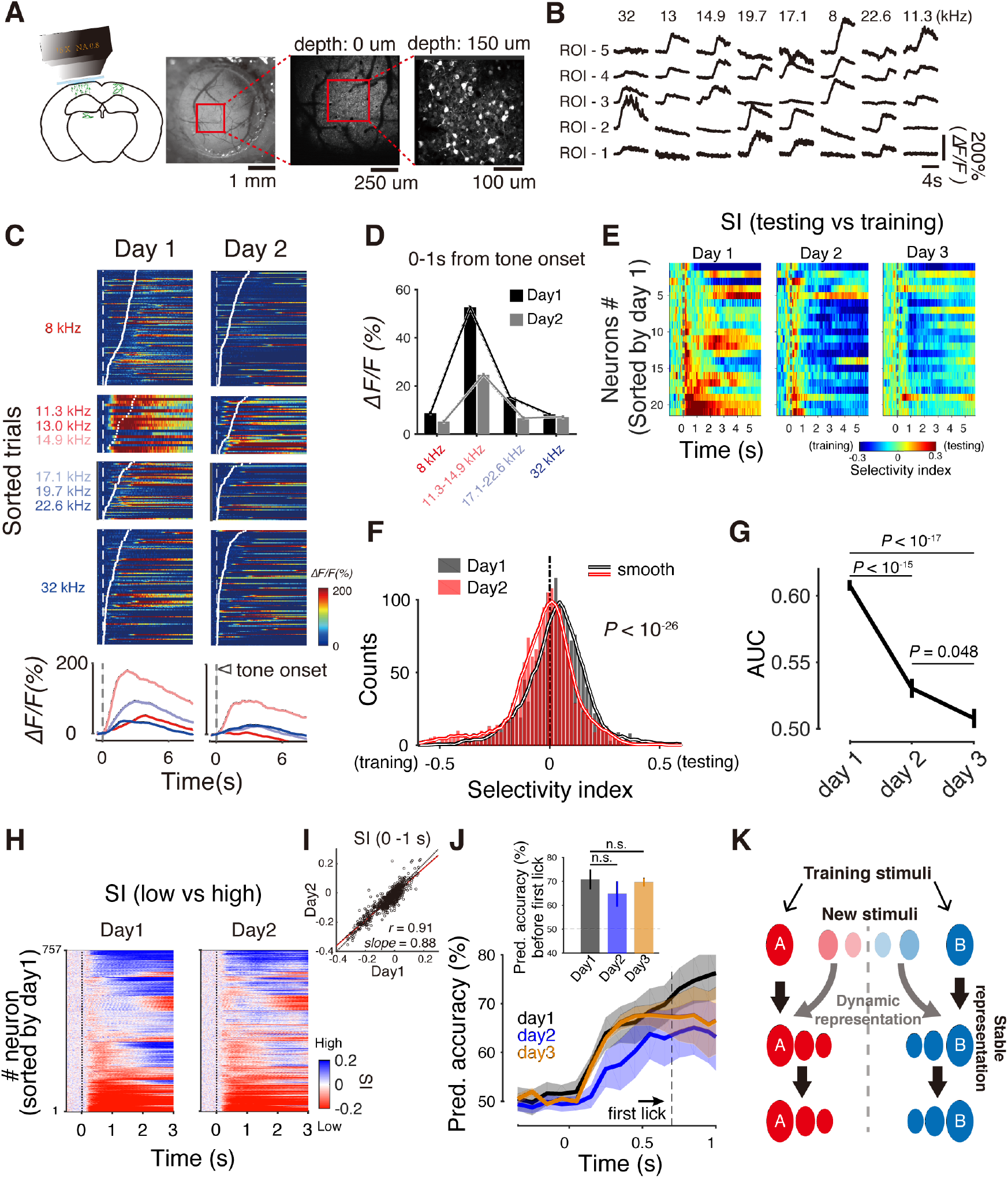
PPC representations show characteristic dynamics of category learning. **(A)** Schematic and images showing in vivo two-photon imaging from PPC populations using GCaMP6s. The same imaging fields were repeatedly imaged across days. **(B)** Example Ca^2^+ signal traces of PPC neurons showing high signal to noise ratio with diverse selectivity and rich dynamics. **(C)** Ca^2^+ signal of an example PPC neuron imaged during behavioral sessions on consecutive days. Upper, color indicates % *ΔF/F*_0_. Blocks of trials presenting training and test stimuli of low and high frequency categories were segregated. Trials within each block were aligned by stimulus onset time and sorted by time of first answer lick. Lower, mean Ca^2^+ signal traces averaged from blocks of trials in the upper panel. Color corresponding to tone frequencies. **(D)** Mean response amplitudes in 1 s window after stimulus onset averaged from different blocks shown in (C). Error bars, s.e.m. **(E)** Selectivity index (SI) between training and test stimuli across time based on ROC analysis using the neuronal responses as shown in c, for all neurons in an example imaging field. Positive values of SI represent preference to test stimuli and negative value for training stimuli. **(F)** Histogram showing distributions of SI values (mean value in 1 s window after stimulus onset) for training vs. test stimuli from the total of 757 neurons across two consecutive days. Significant preference to test stimuli on day 1 (*P* < 10-^23^, Wilcoxon signed rank test), but to training stimuli on day 2 (*P* < 10^−6^, Wilcoxon signed rank test). **(G)** Averaged area under ROC curve, discriminating training and test stimuli, across consecutive days for neurons with significant preference to test stimuli on day 1. Error bars, s.e.m. **(H)** Color raster plot showing index of selectivity between high and low frequency categories across trial time, for all neurons as in **(F)** across consecutive days. Neurons are sorted according to SI values on day 1. **(I)** Scatter plot of the mean SI values in the 1 s trial time window after stimulus onset as in (H) r, correlation coefficient. *Slope*, slope of the linear regression line (red). **(J)** Decoding accuracy across trial time for high and low training tones by population activity of simultaneously imaged neurons in each imaging field using a linear classifier. The classifier was trained using neuronal activity in 80% of training trials on day 1 to decode stimulus categories on consecutive 3 days. Insert, mean decoding accuracies in a 1 sec time window following stimulus onset. **(K)** Schematic showing a conceptual model for differential dynamic representations in PPC reflecting category learning.

**Fig. 3.**
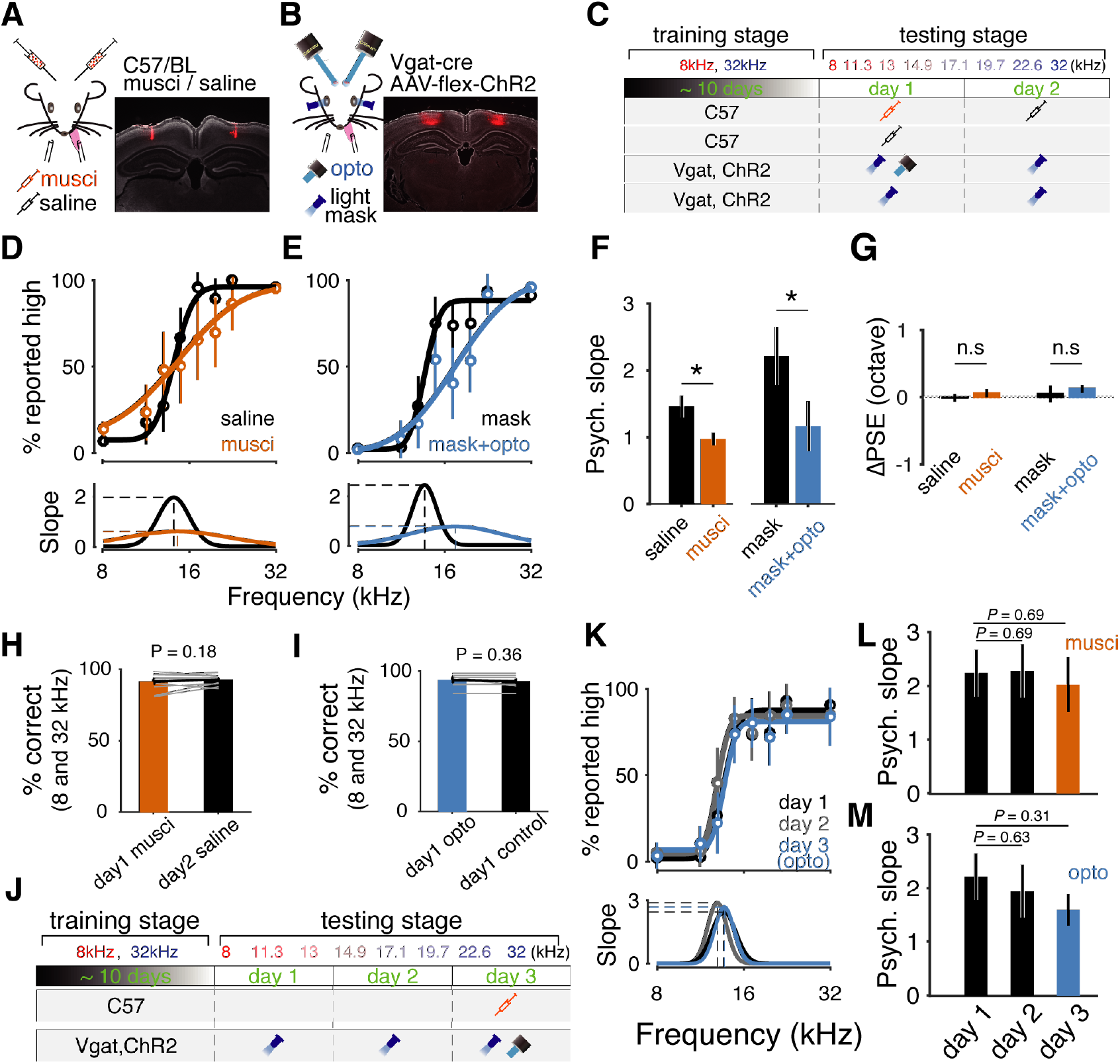
PPC activity is necessary for categorical decision-making on new sensory stimuli. **(A)** Schematic and histology image showing the locations of muscimol or saline injection in PPC. Red color in the histology image is the fluorescence of co-injected red beads to mark the injection sites. **(B)** Schematic and histology image showing the locations of photo stimulation and viral expression of ChR2-mCherry in GABAergic neurons in bilateral PPC of Vgat-Cre mice. Red fluorescence represents mCherry expression. **(C)**, Schematic showing experimental schedules for PPC inactivation and control experiments on the first day introducing test stimuli. Symbols represent the 4 types of experiments as indicated in (A) and (B). Frequencies of tone stimulation at different stages were indicated. **(D)** Psychometric functions (upper) and corresponding slopes (lower) from an example muscimol injection session and control session as indicated in (A) and (C). Error bars, 95% confidence interval. See fig. S9 for individual cases. **(E)** Similar as in (D) for an photoinhibition and control sessions as indicated in (B) and (C). See fig. S10 for individual cases. **(F)** Summary of grouped data of peak psychometric slopes from experiments indicated in (C). Muscimol, *n* = 13, vs. saline, *n* = 14, *P* < 0.05; photoinhibition, *n* = 7, vs. mask only, *n* = 6, *P* < 0.05. Error bars, s.e.m. **(G)** Summary of PSE for experiments indicated in (C) (muscimol vs. saline, *P* = 0.68; photoinhibition with mask vs. mask only, *P* = 0.73; Error bars, s.e.m.). **(H)** Summary of mice’s performance (percent correct) for training stimuli, compared between muscimol injection and control experiments. (*I*) Similar as in (H) for photoinhibition vs. control experiments. Error bars, s.e.m. Paired ŕ-test. (*J*) Experimental schedules, similar as in (C), for PPC inactivation on the third day after introducing test stimuli. **(K)** Similar plot as in e for photoinhibition experiments indicated in (J). See fig. S11 for individual cases. **(L)** and **(M)**, Summary of grouped data of peak psychometric slopes for experiments indicated in (K).

Here we ask whether the PPC neuronal dynamics exhibits characteristics of category learning. With the progress of learning, new sensory stimuli are classified into existing categories, such that the perceptual distinction between new sensory stimuli and exemplars of the same category tend to diminish. We thus focused on whether PPC neuronal activity could distinguish the test stimuli (new sensory information) and the training stimuli (prior knowledge) of the same category, and how such distinction changes with learning. The high performance of perceptual categorization in mice for the test stimuli on day 1 (the first session introducing test stimuli; **Fig. 1**) suggests that significant category learning likely occurred rapidly. Indeed, we found rapid changes in PPC neuronal responses to test stimuli across the first two sessions (days) following introducing test stimuli. On day 1, a significant fraction of neurons responded more strongly to test stimuli than to training stimuli, however, such difference was markedly reduced on day 2 (**Fig. 2, C and D**; also see **fig. S4**).

To quantify the distinction in neuronal responses between test stimuli and training stimuli of the same category, we use receiver operator characteristic (ROC) analysis to define a selectivity index (SI), with negative SI indicating stronger preference to training trials, and positive SI, stronger preference to testing trials (see methods in supplementary materials) (*33*). For learning, the activity in the entire trial is likely to be important, we therefore computed the SI of each neuron in an imaging field for every imaging frame across the time course of the trial (**Fig. 2E**). We found that on day 1, when the test stimuli were first introduced, more neurons significantly distinguish test stimuli and training stimuli, with stronger preference to test stimuli. However, on day 2 and day 3, the preference to test stimuli were largely reduced. This trend is also evident over the entire neuronal population (757 neurons from 6 mice), where the population distribution of SIs also shifted from significantly positive values on day 1 (P<0.001, signed Wilcoxon rank sum test) toward zero on day 2 (**Fig. 2F**). For neurons showing significant preference to test stimuli on day 1, the neuronal discrimination (measured by the area under ROC curve) between test stimuli and training stimuli of the same category also significantly decreased in day 2 and 3 (**Fig. 2G**), indicating that the distinction between the test stimuli and training stimuli of the same category was largely reduced. These results indicate that in parallel to perceptual categorization, PPC neuronal representations exhibit the characteristic dynamics of category learning, incorporating new sensory stimuli into existing categories.

During learning, the circuits not only need to dynamically integrate new information but also need to maintain a stable representation of prior knowledge. We thus ask whether the representation of established “high” and “low” categories in PPC population is relatively stable. First, we found that the number of neurons selective to training stimuli was almost unchanged (**fig. S5E**; 21% in day 1 vs. 20% in day 2). Second, using ROC analysis, we computed the selectivity index of each neuron for “high” vs. “low” frequency categories for all the neurons (757 neurons) over the time course of the behavioral trial across consecutive days (**Fig. 2H**; see methods in supplementary materials). We sort all the neurons according to their SI values on day 1. We found that across the entire population and trial time course, neuronal selectivity for “high” and “low” categories is highly preserved across days, as indicated by the resemblance of the profiles of the SIs between the two consecutive days (**Fig. 2H**; Pearson’s *r* = 0.83). The mean SI values within the time epoch of stimulus and choice (1 s time window after stimulus onset) are also strongly correlated between two consecutive days (**Fig. 2I**; *r* = 0.91). Thus, the neuronal selectivity for established stimulus categories is stable across days. Third, we asked whether the population coding for the established categories was also stable. We used a linear classifier (*34*) to examine the population decoding accuracy of simultaneously imaged neurons in each imaging field for categorical discrimination over the trial time. We used the population activity in 80% of training trials on day 1 as training set to predict either the rest of training trials in day 1, or the training trials in day 2 and day 3 respectively (**Fig. 2J**). We found that the decoding accuracies around the decision time (first lick) were not significantly different for day 1, day 2, and day 3 (**Fig. 2J**), indicating that the population coding for the stimulus categories was stable across days over category learning. Taken together, these results show that PPC neurons exhibit characteristic dynamics of category learning, adaptively classifying new sensory stimuli into existing categories, while maintaining a stable representation for prior learned categories (**Fig. 2K**).

## PPC activity is necessary for perceptual decision-making during category learning

Next, we sought to determine whether PPC activity is important in the decision process during category learning. Our imaging and behavioral data suggest that the learning occurred primarily when the test stimuli were initially introduced in day 1, therefore, it is likely that PPC activity may play a critical role during this period. To test this, we reversibly silenced PPC by bilateral microinjection of muscimol, a selective GABA_A_ receptor agonist, when the new test stimuli were first introduced (**Fig. 3, A and C**). In contrast to previous reports in rodents (*17, 22, 26, 28*) and primates (*23, 24*), we found that mice’s decisions on newly introduced test stimuli were markedly impaired, comparing to control mice microinjected with saline also on day 1. The impairment is measured as a significant reduction in the peak discriminability (slope at the category boundary; saline, 1.46 ± 0.16; muscimol, 0.97 ± 0.09; *P* < 0.05, Wilcoxon rank sum test; **Fig. 3, D and F**).

Since muscimol inactivation does not have the temporal resolution to distinguish whether it was during training trials or during testing trials that silencing PPC led to impaired categorization, we employed optogenetics to inactivate PPC in a more temporally controlled fashion (25). An adeno-associated virus (AAV) encoding Cre-inducible channelrhodopsin 2 (ChR2) (*35*) was injected bilaterally in PPC of VGAT-Cre mice (*36*) to express ChR2 in GABAergic inhibitory neurons. Glass optical windows were implanted above PPC of both hemispheres following virus injection. Collimated blue laser beams were used to illuminate the glass window to activate ChR2 in GABAergic neurons to inhibit the illuminated cortical areas (**Fig. 3, B and C**; also see **Movie S1**) (*25*). During task performance, photoinhibition was applied in all testing trials, but only in 10% of training trials for comparison, leaving majority (90%) of training trials unaffected. We found that this manipulation also significantly reduced mice’s categorization performance for the new tone stimuli (control, 2.43 ± 0.46; photoinhibition, 1.17 ± 0.37; *P* < 0.05, Wilcoxon rank sum test; **Fig. 3, E and F**), indicating that PPC activity during testing trials rather than during training trials is required for categorical decisions on the new test stimuli. Moreover, we found that when photoinhibition of PPC was applied in 50% of the testing trials, mice’s categorization performance was not significantly different in photostimulated trials comparing to that in control trials (**fig. 7**), suggesting that PPC activity in partial of the testing trials on day 1 was sufficient for category learning of the newly introduced tone stimuli.

These results, together with our imaging data (**Fig. 2**), point to the possibility that PPC activity is specifically required only during category learning for new stimuli, but not after learning. Consistent with this idea, for the training stimuli, neither muscimol silencing (**Fig. 3H**), nor photoinhibition (**Fig. 3I**) influenced mice’s performance (also see **fig. 8**). To further test this possibility, we performed bilateral PPC perturbation in a separate cohort of mice after the test stimuli had been well-experienced following continued training for two additional sessions (>1000 trials; **Fig. 3J**). We found that when the test stimuli were well-experienced, inactivation of PPC did not significantly influence mice’s categorization performance (**Fig. 3, K** to **M**), agreeing with previous studies in rodents and non-human primates assessing the effect of PPC inactivation on choice behavior, where the stimuli were likely also well-learned (*17, 23, 25, 26*). Taken together, these data indicate that PPC is causally involved in categorical decisions specifically on new sensory stimuli during category learning, but not after learning when the stimuli have been already incorporated in to existing categories.

## PPC activity counterbalances biases from recent trial history

Decision-making in humans and animals is often biased by recent behavioral history, which usually produce a negative impact on performance in complex and unpredictable settings (*21, 22, 37–41*). It has been reported that PPC activity reflected both accumulated behavioral history (*15, 16, 21, 22*) and systematic bias from long-term training history (*8*). Meanwhile, our imaging data, as well as previous studies (*6, 11, 15, 24*), show that even after learning, PPC neurons continue to encode stimulus category and choice information. To understand how the accumulated prior knowledge encoded in PPC may influence the bias from recent behavioral history, we examined the effect of PPC inactivation on trial-by-trial history bias during the auditory categorization task. We quantified the influence of the preceding training trials on choices for testing stimuli by comparing mice’s performance on test stimuli when the stimulus in the immediate preceding training trial was of the same vs. different category as the current test stimuli. Under control condition, preceding training trials did not significantly bias mice’s performance on the testing trials (**Fig. 4B**, 77.3 ± 4.6% for same category vs. 74.3 ± 3.2% for different category, P = 0.32), consistent with a previous study, in cases where decisions did not depend on short-term memory (*22*). However, when PPC was inactivated by either muscimol or photoinhibition, preceding training trials significantly biased mice’s choices on test stimuli *towards* the preceding categories (**Fig. 4A**, muscimol, same 71.8 ± 3.4% vs. different 63.5 ± 3.7%, P = 0.017; photoinhibition, same 70.3 ± 2.4% vs. different 55.9±3.2%, P = 0.003.) Thus, whereas mice’s decisions were minimally influenced by preceding events when PPC was intact, inactivation of PPC led to a significant bias by short-term behavioral history (**Fig. 4, C and D**). Moreover, when the newly introduced test stimuli became well-experienced after continued training, inactivation of PPC no longer resulted in significant bias by preceding training trials (**fig. S12, E and F**). These results indicate that PPC activity counterbalances the short-term history bias when categorizing novel sensory stimuli.

**Fig. 4.**
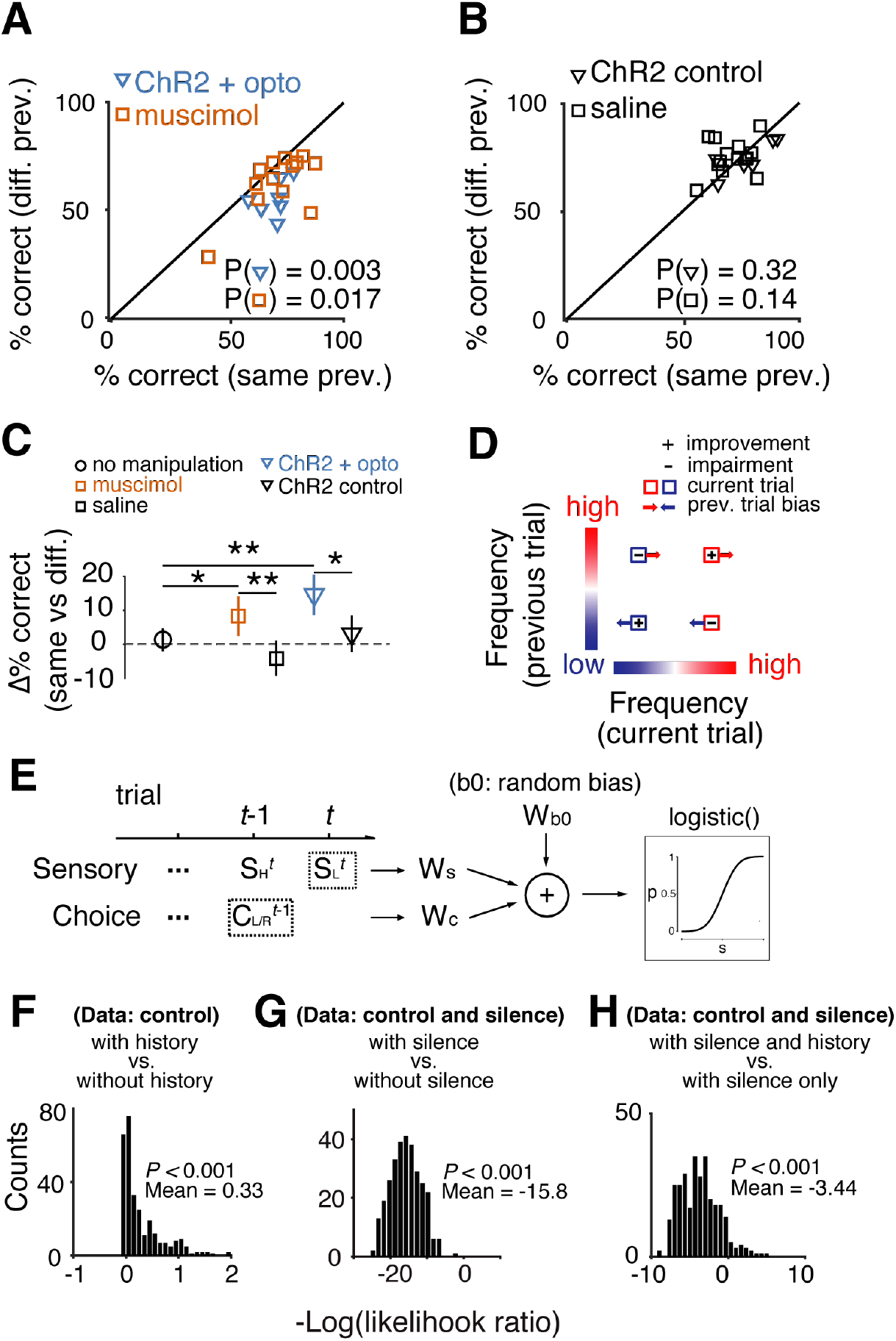
PPC activity counterbalances short-term history bias on perceptual decisions. **(A)** Comparison of behavioral performance on testing trials when the stimuli in preceding training trial was of different categories (diff. prev.) or the same category (same prev.) as the current trial. Mice’s choices were significantly biased by the previous trial when PPC was silenced by muscimol (P = 0.017, *n* = 13) or by photoinhibition (P = 0.003, *n* = 7). **(B)** Same plot as in (A). Mice’s choices were not significantly biased by previous trials under control conditions (saline, *P* = 0.14, *n* = 14; light mask, *P* = 0.32, *n* = 6). **(C)** Summary of changes in performance attributable to previous trials as in (A) and (B). Data from a separate group of control mice without any manipulation were also included. *, *P* < 0.05; **, *P* < 0.01. Two sample *t*-test. Error bars, s.e.m. **(D)** Schematic showing the bias effect on categorical decisions by the previous trials when PPC was silenced. **(E)** Illustration of linear regression models to simultaneously quantify the effects of PPC silencing on categorization slopes and short-term history bias. **(F)** to **(H)**, Likelihood ratio test comparing different regression models using data from the control group for models with or without previous trial choices **(F)**, using all data from both control and PPC silencing groups for models with or without the interaction term between sensory stimuli and PPC silencing **(G)**, and using data from both groups for models with or without the interaction term between PPC silencing and previous choices (H).

To simultaneously examine PPC’s contributions to category learning (**Fig. 3**) and to short-term history bias (**Fig. 4, A and C**), we modeled animals’ performance in control and silencing conditions using a series of logistic regression models (Generalized Linear Mixed-Model, GLMM; **Fig. 4E**; also see methods in supplementary materials) (*39*). Our models confirmed the lack of history bias effect in control data, in that including of previous choice as a regressor term did not significantly improved the model prediction of the behavioral data (**Fig. 4F**). The models also confirmed the effect of PPC inactivation on changing the psychometric slopes, in that including of PPC silencing as a regressor significantly improved the model prediction (**Fig. 4G**; also see **fig. S13, A to D**). Interestingly, when including an interaction term between previous choice and the silencing condition as a regressor, the model predicted animals’ behavior significantly better comparing to the model that only quantified the effect of silencing on the psychometric slope changes (**Fig. 4H**; see **fig. S14, A to C**), indicating that the effect of PPC silencing on short-term history bias is in addition to its effect on categorization slopes. Thus, PPC causally contributes to the decision process in category learning by both influencing categorization slopes and counterbalancing short-term history bias when categorizing novel stimuli.

## PPC to auditory cortex projections are important for categorical auditory decisions

PPC neurons consist of diverse populations interconnected with many brain regions (*42*), and are involved in many behavioral functions (*6, 7, 9, 14, 18, 43*). It is likely that for a given task, specific circuits interconnecting PPC and task-specific brain regions are critically involved. We thus further examined the circuit-level specificity of the causal role of PPC neurons in auditory categorical decision-making by manipulating PPC to auditory cortex projections using chemogenetics. We first determined the projection patterns between PPC and the auditory cortex by expressing green fluorescent proteins (GFP) in PPC, and red fluorescent proteins (tdTomato) in the primary auditory cortex (AUDp), using AAV vectors. We found that PPC receives projections from AUDp while sending projections to the dorsal portion of auditory cortex (AUDd; **Fig. 5A**), implying an interconnected circuit between PPC and the auditory cortex (**Fig. 5B**). We next expressed hM4Di, a designer receptor exclusively activated by designer drug (DREADD), in PPC of both hemispheres using an AAV vector (AAV-Syn-hM4Di). On day 1 (test tones first introduced), we injected Clozapine-N-oxide (CNO) to bilateral AUDd to silence the projections from PPC prior the behavioral session (**Fig. 5C**). As control, in a separate group of mice we injected AAV-Syn-mCherry in PPC, and then injected CNO to AUDd during task performance on day 1. We found that comparing to the control group, silencing PPC to AUDd projections impaired mice’s categorization performance for the new tone stimuli, as indicated by a significant reduction in the slopes of psychometric functions (**Fig. 5D**). Quantification using GLMM test also shows a significantly better prediction of the model with the term of projection silencing than without the term (**Fig. 5E**). These results indicate that the specific projections from PPC to secondary auditory cortex play a critical role in the auditory categorical decisions on new sensory stimuli.

**Fig. 5.**
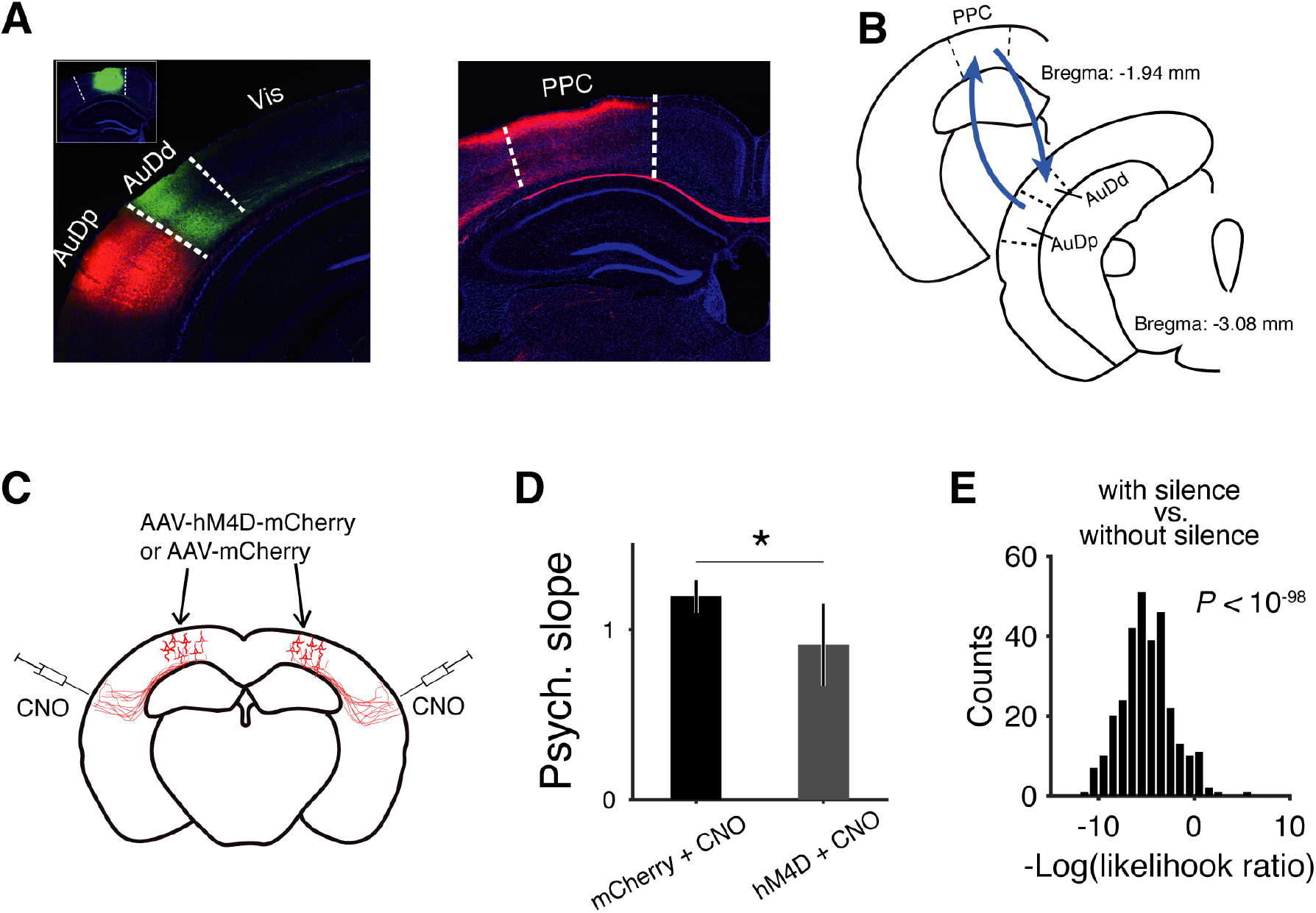
PPC to auditory cortex projections are necessary for categorical decision-making on new sensory stimuli. **(A)** Histology images showing the projection patterns between PPC and the auditory cortex. Left, red fluorescence indicates the site of injection of AAV-syn-mCherry in the primary auditory cortex. Green fluorescence in the dorsal auditory cortex indicates axons projected from PPC. Insert, showing site of injection of AAV-syn-GFP in PPC. Right, red fluorescence indicates axons projected from the primary auditory cortex in PPC. **(B)** Schematic showing the projection patterns between PPC and the auditory cortex. **(C)** Schematic showing chemogenetic silencing of the projections from PPC to the auditory cortex. AAV-syn-hM4D-mCherry (experimental group) or AAV-syn-mCherry (control group) were injected in bilateral PPC. CNO was microinjected in bilateral auditory cortex prior the first session introducing test tone stimuli. **(D)** Summary of the effect of silencing PPC to auditory cortex projections on psychometric slopes. *P* < 0.05, Wilcoxon rank sum test, *n* (control) = 7, *n* (silencing) = 8. Error bars, s.e.m. **(E)** Likelihood ratio test comparing regression models with or without projection silencing as a regressor.

The neural basis for how organisms integrate novel sensory information with prior knowledge to assign behavioral meanings to physical world remains poorly understood. Here we provide functional and causal evidence to support a critical role of PPC circuits in this process. We show that PPC neurons exhibit characteristic dynamics related to category learning, adaptively incorporating new sensory information while maintaining stable representations for learned category to choice mappings, a balance in flexibility and stability. Importantly, we provide, to our knowledge, the first evidence supporting that PPC causally contributes to perceptual decision-making in categorizing new sensory information, a key process in category leaning. Furthermore, at the circuit level, we identified a necessary role of PPC to auditory cortex feedback projections, providing a critical foundation for further dissection of circuit-level mechanisms of category learning. After the stimuli were well-learned, PPC continued to represent stimulus categories (**Fig. 2**) and behavioral choices (*6, 11, 13, 23, 24*). This is likely to be important for guiding future decisions on new sensory stimuli, and reducing biasing effects from random factors (**Fig. 3**). Some of recent studies showed that silencing PPC impaired visually guided (*14, 17, 27, 28*) but not auditory guided (*17, 26, 28*) choice behaviors in rodents, while in primates, silencing parietal cortex did not influence either visual motion (*23*) or self motion (*24*) guided choice behavior, suggesting that that PPC may also play certain visual-specific functions in rodents. Our results, however, support a more general functional role of PPC in perceptual decision-making when facing novel conditions. More recent studies in rodents also using well-trained stimuli showed increased performance in tasks requiring short-term memory or working memory following inactivation of PPC (*21, 22*), likely attributable to an interference with short-term memory by the effect of PPC on recent choice history, but not an direct effect on perceptual decision-making *per se.* This is consistent with our observation that PPC activity counterbalances recent history bias. Our results thus provide a framework for mechanistic understanding of neuronal circuit basis of decision-making integrating new sensory information with prior knowledge in a dynamic environment.

## Author contributions

L.Z. and N.L.X. conceived the project and designed the experiments. L.Z. performed all the experiments and data analysis. Y.Z. and L.Z. performed the chemogenetic experiments. L.Z. and C.A.D. conceived and performed the GLMM analysis. L.Z., J.P. and N.L.X. developed the hardware and software of the behavior and imaging system. L.Z. and N.L.X. wrote the manuscript with contributions from C.A.D.

## Supplementary Materials and Methods

### Materials and Methods

#### Animal subjects and behavior

Experimental procedures were approved by the Animal Care and Use Committee of the Institute of Neuroscience, Chinese Academy of Sciences. Data were acquired from male C57BL/6J (SLAC), and VGAT-Cre mice (Jackson Laboratory, JAX 017535) (*36*), age 8-10 weeks at the start. Mice used for behavioral tests were housed in a room with a reverse light/dark cycle. Mice had no previous history of any other experiments. On days not tested, mice received 1 ml of free water. On days of testing behavioral sessions last for 1 to 2 h where they received water mainly from task performance and with supplements to reach 1 *ml.* Each mouse’s weight was measured daily to ensure that weight loss did not exceed 20 % of the mouse’s pre-water restriction weight.

After ~7 days’ recovery following surgery, mice were started with water restriction procedure. Each mouse received 1 ml water per day and the body weight was monitored. After ~7 days of water restriction, behavior training was started. (*32, 44*). Water consumption was calculated as total water = number of rewarded trials × ~ 6 *μl* water in one delivery for each training session. If mice consumed less than 0.5 ml water, additional water supplement was provided. Mice were allowed to perform the task until sated.

The behavior was divided into training stage and testing stage. In the training stage, mice learned to discriminate tones of two easy frequencies, separated by 2 octaves (8 kHz and 32 kHz). Each trial was started with a 500-1000ms random delay, followed by tone stimuli (duration, 300 ms). Mice were required to respond with licking left or right lickport within a 1~3 s answer period to indicate their choices. Correct answers were defined as licking the left lickport in response to the lower frequency tone (e.g., 8 kHz), and licking the right lickport in response to the higher frequency tone (e.g., 32 kHz). Correct responses lead to the water valve open to dispense a small amount of water reward (~6 ul). Error responses lead to a 2~4 s time-out punishment, during which licking to the wrong side reinitiates the time-out period. If mice made no response lick within the 3 s response window, the trial was defined as a “miss” trial, leading to the initiation of inter-trial-interval (ITI). Mice reached criteria of >85 % correct in 7-10 days (**fig. S1**).

During testing stage, 8 different tones (in frequency: 8000, 11300, 13000, 14900, 17100, 19700, 22600 and 32000; in octave difference related to 8000 Hz: 0, 0.5, 0.7, 0.9, 1.1, 1.3, 1.5 and 2) were presented in pseudorandomly interleaved trials. Trials containing newly introduced frequencies were defined as “testing” trials, constituting ~30% of total trials. The psychometric functions were fitted to choice data using a 4-parameter sigmoidal function (*45*):

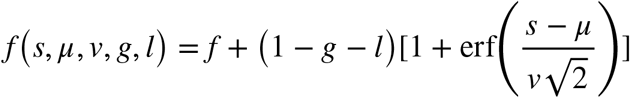

where *s* is the tone frequency in octave, *μ* is the mean value of the distribution representing subject’s point of subjective equality (PSE), *ν* is the variation of the distribution representing the subjects discrimination sensitivity. *g* and *l* are the guess rate and lapse rate.

#### Behavioral system

Experiments were conducted inside custom-designed and fabricated double-walled sound-attenuating boxes. Mice were head-fixed with a pair of clamps and thumbscrews. The mouse body is contained in an acrylic ‘body tube’ (1 inch i.d.), with the mouse head extending out and the front paws gripping the tube edge or a ledge after head-fixation (**Movie S1**). The holder and body tube in turn are attached to a custom-designed tube holder mounted to a caddy which is mounted to the behavior box with table clamps after head-fixation. Water rewards are provided by two custom-made metal lickports placed on the left and right side of the mouse mouth. The lickports are connected to a custom-desgined capacitive-sensing circuit board that reliably detects the contact of the tongue during licking. The amount of water delivered was controlled by the open time a solenoid water valve calibrated at least once a week. Calibration was achieved by estimating the time of valve opening necessary to deliver the desired volume per trial, by running 100 deliveries and calculating water volume by weight. The fluid pressure was not adjusted within each behavioral session.

The mouse auditory discrimination behavior was controlled by a custom-developed low-cost and high-precision real-time control system, the PX-Behavior System (**fig. S1**). The system includes a custom-designed tone-generating module (TGM) to generate sound waveforms of arbitrarily high frequencies with high fidelity, and a microcontroller (arduino MEGA 2560) implementing a real-time virtual state machine framework for programming the behavioral protocols, stimulus delivery, and measurement of behavioral events. The arduino board communicates with the TGM through a custom-designed cache board without interrupting the flow of the behavioral protocols. The TGM sends the specified sound waveforms to an amplifier (ZB1PS, Tucker-Davis Technologies) to drive a speaker (ES1, Tucker-Davis Technologies) to produce acoustic stimuli. 5 ms cosine ramps are applied to the rise and fall of all tones. Behavioral data were logged via serial port using custom-written software in python.

The sound system was calibrated using a free-field microphone (Type 4939, Brüel and Kjær) over 3–60 kHz and showed a smooth spectrum (± 5 dB). Measurements were performed with the behavioral box closed and the microphone positioned at the location and orientation of the mouse ear-position in the presence of the mouse mounting system. The microphone was connected to a preamplifier (Type 2670, Brüel and Kjær), and signals were digitized with a National Instruments acquisition card (NI 9201) at 500,000 samples per second for analysis with custom software developed in LabVIEW (National Instruments).

#### Surgery and virus injection

During surgery, mice were anaesthetized with isoflurane (~2%). For inactivation experiments, we targeted bilateral PPC (2 mm caudal, 1.5 mm lateral to bregma; both hemisphere) by microinjection of muscimol in wild-type mice, or viral solution in VGAT-Cre mice. For photoinhibition experiments, craniotomies were made over the PPC of VGAT-Cre mice following virus injection. Viral solution containing AAV2/8-CAG-FLEX-ChR2-mCherry (titer: ~10^13^ genomes/mL), or AAV2/9-Syn-FLEX-ChrimsonR-tdTomato (titer: ~10^13^ genomes/mL; Shanghai Taitool Bioscience Co.Ltd) was injected slowly (~ 100nL over ~20 minutes). The injection system comprised of a pulled glass pipette (25–30 μm O.D. at the tip; Drummond Scientific, Wiretrol II Capillary Microdispenser) back-filled with mineral oil. A fitted plunger was inserted into the pipette and advanced to displace the contents using a hydraulic manipulator (Narashige, M0-10). Retraction of the plunger was used to load the pipette with virus solution. The injection pipette was advanced into the brain using a Sutter MP-225 micromanipulator. After injection, the craniotomies were covered with a circular glass coverslip, sealed in place with dental cement (Jet Repair Acrylic, Lang Dental Manufacturing). For pharmacological inactivation, the locations of PPC of both hemispheres were marked for later injection. Mice were allowed at least 7 days to recover before water restriction.

For chronic two-photon in vivo imaging, a circular craniotomy (2-3 mm diameter) was made over the PPC of the left hemisphere, followed by injection of AAV2/9-hSyn-GCaMP6s (titer: ~10^13^ genomes/mL; Shanghai Taitool Bioscience Co.Ltd) at a depth of ~250 μm. After injection, a double-layered imaging window was implanted at the craniotomy as described previously (*32*). The double-layered glass comprised two 200-μm-thick glass coverslip (diameter, ~3 mm) attached to a larger glass coverslip (diameter, 5 mm) using ultraviolet cured optical adhesives (Edmund Optics). Following virus injection and window implantation, a titanium head-plate was then attached to the skull with cyanoacrylate glue and dental cement to permit head fixation during behavior.

#### Pharmacological inactivation of PPC

We used the GABA (c-aminobutyric acid) agonist muscimol hydrobromide (Sigma-Aldrich, dissolved in saline, 5 mg/ml) to reversibly silence PPC by stereotactic injection. We used the same nano-injection system used in our virus injection to deliver a small volume with slow injection speed (50 nL at 10 nL per min) to ensure accurate and localized injection volumes (*32*), which minimized the spread of muscimol to neighboring brain regions comparing to conventional cannula based infusion system. A diluted (1:20) red beads was added to identify the injection sites for both muscimol and saline injections.

#### Photoinhibition

For optogenetic inactivation of PPC, blue laser (473 nm) was delivered to the PPC of both hemisphere through two collimators connected to a splitting fiber optics coupled with a diode-pumped solid state laser (DPSS; Shanghai Laser & Optics Century Co., Ltd.). Photostimulation comprised of trains of pulses at 40 Hz with pulse width of 5 ms. Laser power at the cranial windows (beam size ~1 mm in diameter) was calibrated before each session start (6.5-8 mW/ mm2). To prevent the mice from disturbance by blue light illumination, a ‘masking flash’ light was delivered with the same pattern of photostimulation throughout each trial using two blue LEDs positioned near the eyes of the mice (**Movie S1**).

We examined whether manipulation of PPC activity directly influenced mice’s motor output by plotting reaction time and licking (**fig. S14**). We observed no significant changes in mice’s reaction time when PPC was silenced by either muscimol or photoinhibition comparing to control mice (**fig. S14, A and B**). Neither did we observed significant changes in mice’s lick rate during response period when PPC was silenced (**fig. S14, C and D**).

#### Chemogenetic inactivation of PPC projections

We used a DREADDs based method to silence the PPC to dorsal auditory cortex projections (*46*). AAV2/9-hSyn-DIO-hM4D(Gi)-mCherry (1.72 × 10^12^ genomic copies per mL) was injected to bilateral PPC, and was allowed for 4 weeks of expression. Clozapine-N-Oxide (CNO) was dissolved in saline (0.9% NaCl solution) to a stocking solution of 20 mg/mL stored at −20°C, and was diluted to a working concentration of 1 μg/μl each day before experiments. During the day of experimental testing, saline or CNO (1 μl each hemisphere,1 μg/μl) was microinjected to bilateral AUDd, 30~40 min before behavioral testing.

#### Two-photon in vivo calcium imaging

Imaging was performed using a custom built two-photon microscope (http://openwiki.janelia.org/wiki/display/shareddesigns/MIMMS). GCaMP6s was excited using a Ti-Sapphire laser (Chameleon Ultra II, Coherent) tuned to 920 nm. Images were acquired using a 16x 0.8 NA objective (Nikon), and the GCaMP6s fluorescence was isolated using a bandpass filter (525/50, Semrock), and detected using GaAsP photomultiplier tubes (10770PB-40, Hamamatsu). Horizontal scanning was accomplished using a resonant galvanometer (Thorlabs; 16 kHz line rate, bidirectional). The noise produced by the resonant scanner were attenuated to < 30 dB using an optical window sealing the output opening of the resonant scanner module. The entire microscope was enclosed in a custom-designed and custom-fabricated, double-walled sound attenuation box (internal noise level < 30 dB with the two-photon imaging system running). The average laser power for imaging was ~ 70 mW, measured at the entrance pupil of the objective. The field of view was 300 by 300 um (512 x 512 pixels), imaged at ~30 frames/s. The system was controlled using ScanImage (http://scanimage.org) (*47*). The initiation of each trial was synchronized between image acquisition and behavior by a trigger signal sent from our PX-Behavior System to the ScanImage system. One imaging field was acquired for each of the six mice over 2 days (2 mice) or 3 days (4 mice), after the test tone stimuli were first introduced. The imaging fields in day 2 and 3 were aligned to that in day 1 based on the superficial blood vessels in zoom-out view followed by careful alignment to single neurons at higher level magnification.

#### Imaging data analysis

To correct for brain motion, all imaging frames from each imaging/behavior session were aligned to a target image frame using a cross-correlation-based registration algorithm (discrete fourier transformation, DFT, algorithm) (*48*). The target image was obtained by mean projection of image frames from a trial visually identified to contain still frames. To extract fluorescence signals from individual neurons, regions of interest (ROIs) were drawn manually based on neuronal shapes. Mean, maximum intensity, and standard deviation values of all frames of a session were used to determine the boundaries of the neurons. The pixels in each ROI were averaged to estimate the time series of fluorescence (*F*) of a single cell. Before calculating the Δ*F/F_0_*, slow calcium fluorescence change correction were performed by minus the eighth percentile of ROI fluorescence within a 200s time window for each frame. For each ROI, Δ*F/F_0_* (%) was calculated as (Δ*F/F_0_*)× 100, where *F_0_* is the index of the peak of the histogram of *F*.

To quantify single neuronal selectivity differentiating training stimuli and newly introduced test stimuli, or the “high” and “low” categories, we used an ideal observer decoding based on ROC analysis (*33, 49*). We defined relative selectivity index (SI) as *SI* = 2 × (*AUC* – 0.5)which is a measure with a range from −1 to 1. For selectivity for “training” vs. “testing”, SI < 0 means “preferring training stimuli”, and SI > 0 means “preferring test stimuli” and 0 represents no selectivity. For stimulus category selectivity, SI < 0 for “high” and SI > 0 for “low”. SI across the time course of the behavioral trial is computed from frame by frame Ca^2+^ signals (**Fig. 2, E and H**). For calculating the SI in a specific period, e.g., 0~1s after tone onset, average of Δ*F/F_0_* values in this period in each trial were used.

We used a linear classifier based on support vector machine (SVM) with linear kernels. Simultaneously imaged calcium signals were arranged into a M by N matrix, where M is the number of trials and N is the number of neurons. Each element in the matrix is the Δ*F/F_0_* of a particular neuron at a given image frame in a trial. Such matrices were prepared for all imaging frames across time. To decode the stimulus categories on day 1, we used randomly selected 80% of the training trials on day 1 as the training set to train the linear-kernel SVM classifier, and used the rest 20% of training trials to test the decoder, and repeated this process for 1000 times to obtain the average accuracy of decoding. To decode the stimulus categories on day 2 and 3, we also used data in the training trials on day 1 as the training set, to predict the training trials on day 2 and 3 (**Fig. 2J**).

#### Linear regression models

To address the contribution of multiple variables on animals’ behavior (auditory stimuli in octave, effect of silencing PPC, and previous trial history), we fitted a series of Generalized Linear Mixed-Model (GLMM) using the Matlab function fitglme. First, to characterize the effect of previous trial history in control sessions (**Fig. 4F**), we fit a model where the probability of rightward choice was a logistic function of pure tone stimuli (in octave) and animals’ choice on the previous trial. We compared this model with a simpler model without the previous trial effect (see GLM model comparison), and found that including the previous trial effect resulted in a significantly worse fit to control behavior (*P* < 0.001).

Second, we quantified the effect of silencing bilateral PPC neurons on animals’ psychometric performance (**Fig. 4G**) by including an interaction term between auditory stimuli and the silence condition. The statistics of this interaction term reflect a change in the slope of the psychometric curves (**fig. S12**). Adding this interaction term significantly improved the fit to animals’ behavior compared to the simpler model of fitting auditory stimuli alone (*P* < 0.001).

Finally, we examined whether silencing PPC neurons also changed the effect of previous trial history (**Fig. 4H**), in addition to the silencing effect on psychometric slopes. For this final model, choice was fit as a logistic function of stimuli, the interaction between stimuli and the silencing condition, and the interaction of previous choice and the silencing condition (**fig. S13**). The final model was a significant better fit to behavior than the model without the last interaction term (*P* < 0.001). For all the GLMM models, the rightward and leftward choice was defined as 1 and 0 respectively. For the silence condition, 0 and 1 represent if the current trial was from a control or silence session. Random effects of psychometric biases were included for individual animals.

#### Model comparison

To justify more complex models of behavior, we conducted model comparisons using the cross-validated negative log likelihood measure. We always compared a test model with one extra predictor (e.g., the interaction between stimuli and the silence condition) against a null model without that extra term. For each round of cross-validated model prediction, 70% of the trials were randomly selected as the training set, with the rest 30% as the testing set. The same data set was used for both the test model and the null model. The negative log likelihood ratio (log LR) for that round is the difference between the negative log likelihoods of the two models (test – null). This procedure was repeated for 300 times, and the distribution of –log LR was compared to 0. If the distribution of –log LR was significantly smaller than 0, this means that the test model out-performed the null model in predicting animals’ behavior, thus justifying adding the extra term.

#### Supplementary Text

PPC is considered to be at the interface between sensory and motor system and may represent movement intention (*43*). To control for the effect on motor output, we examined whether manipulation of PPC activity influenced mice’s reaction time or licking (**fig. S14**). We observed no significant changes in mice’s reaction time when PPC was silenced by either muscimol or photoinhibition comparing to control mice (**fig. S14, A and B**). Neither did we observed significant changes in mice’s lick rate during response period when PPC was silenced. The changes in the categorization slope or short-term history bias after PPC silencing were unlikely due to changes in motor output.

**Fig. S1.**
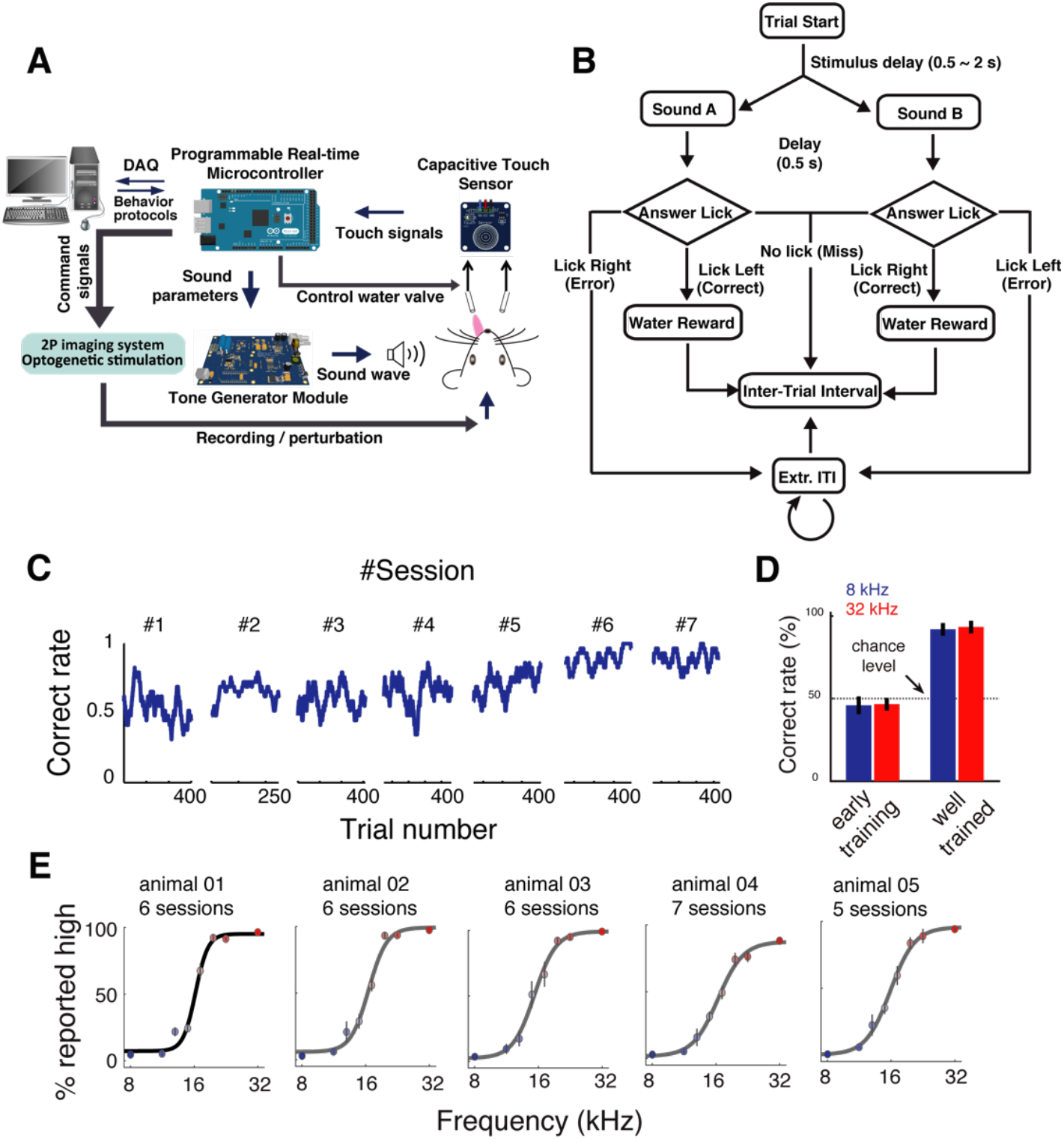
Behavioral control system and auditory psychophysical task in mice. **(A)** Schematic diagram of custom designed behavior control hardware system for two choice auditory decision task in head-fixed mice, named as PX-Behavior System. **(B)** Schematic diagram of the auditory two-alternative forced choice task in head-fixed mice. **(C)** Example of auditory 2AFC task performance over the time course of training. **(D)** Summary of mice performance (percent correct, mean ± s.e.m.) upon training stimuli at the beginning of training and at well-trained stage. **(E)** Psychometric curves from a separate group of mice with randomly rewarded testing trials, i.e., the reward in the trials presenting intermediate tone frequencies were rewarded randomly, not according to the experimentally defined categorization boundary. Data from 5-7 continuous sessions following test stimuli were first introduced were grouped for each animal. Circles with bars, mean ± s.e.m.

**Fig. S2.**
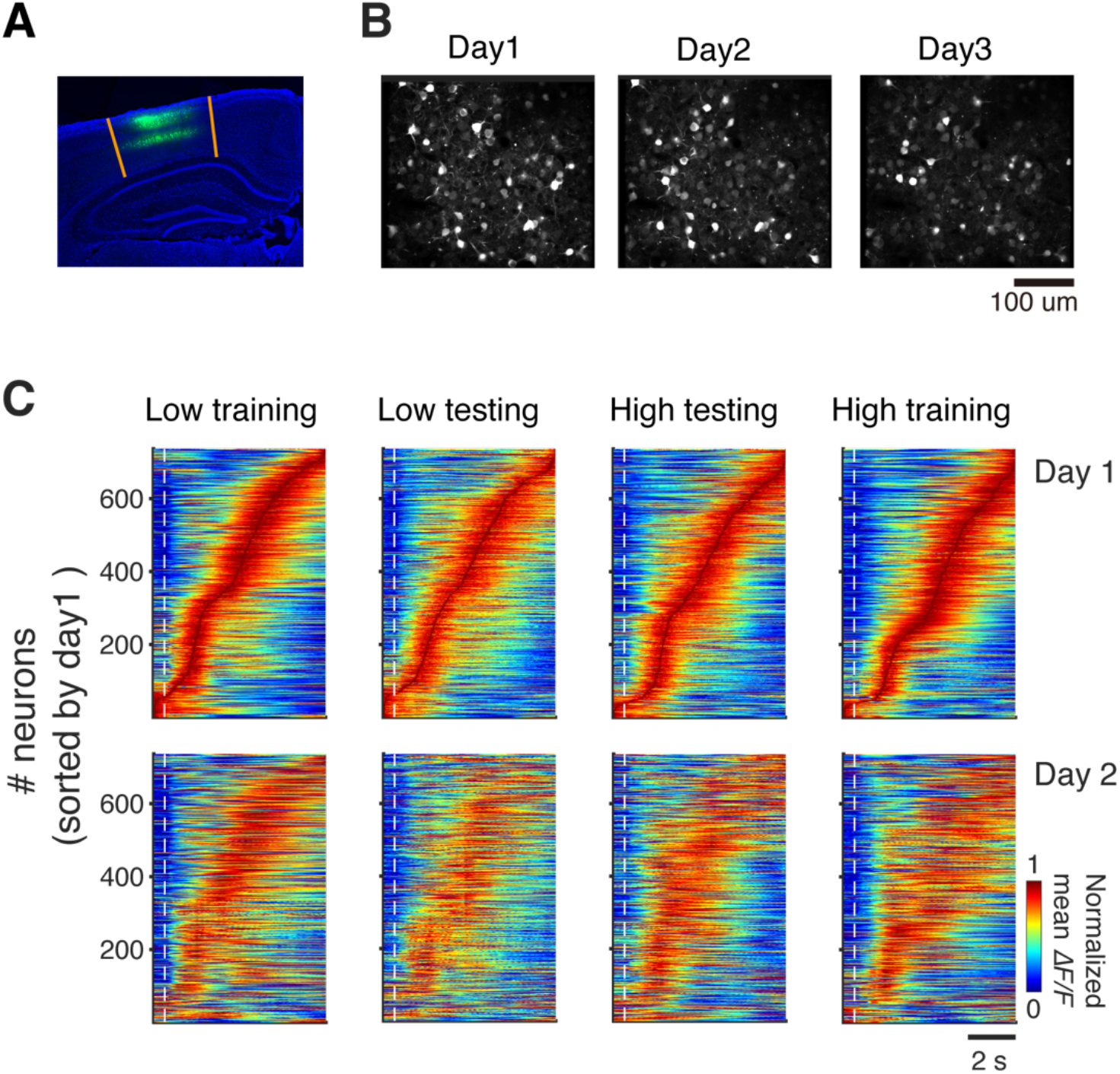
Overview of population response profiles in PPC across consecutive days recorded using two-photon calcium imaging. **(A)** Expression of GCaMP6s in PPC using AAV-SYN-GCaMP6s. **(B)** An example imaging field across three days. **(C)** Normalized mean Ca^2+^ signals from all neurons (736 neurons from 5 mice) in different trial types (training and testing, with “low” or “high” stimulus categories). Neurons were sorted by the response peak time in each trial type on day 1 (test stimuli were first introduced). Upper, data from day 1. Lower, data from day 2. Dash lines indicate stimulus onset time.

**Fig. S3.**
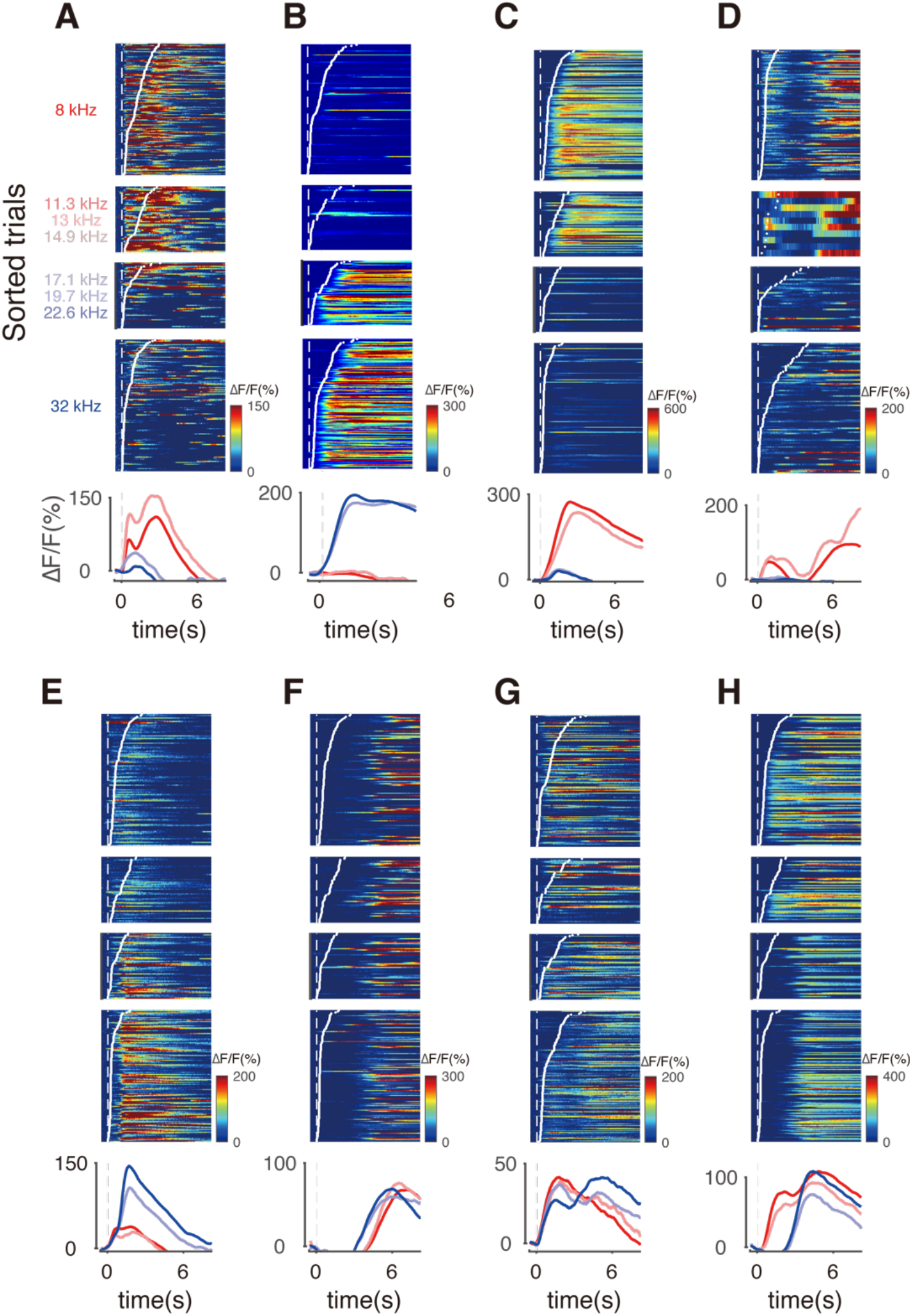
Example neurons showing diverse response patterns during auditory categorization task. Testing trials with the same categorical frequencies were grouped together. Dash lines indicate stimulus onset. Ca^2+^ signals (Δ*F/F*) in each trial were aligned to stimulus onset, and sorted according to first lick time after stimulus onset. Traces are mean across trials.

**Fig. S4.**
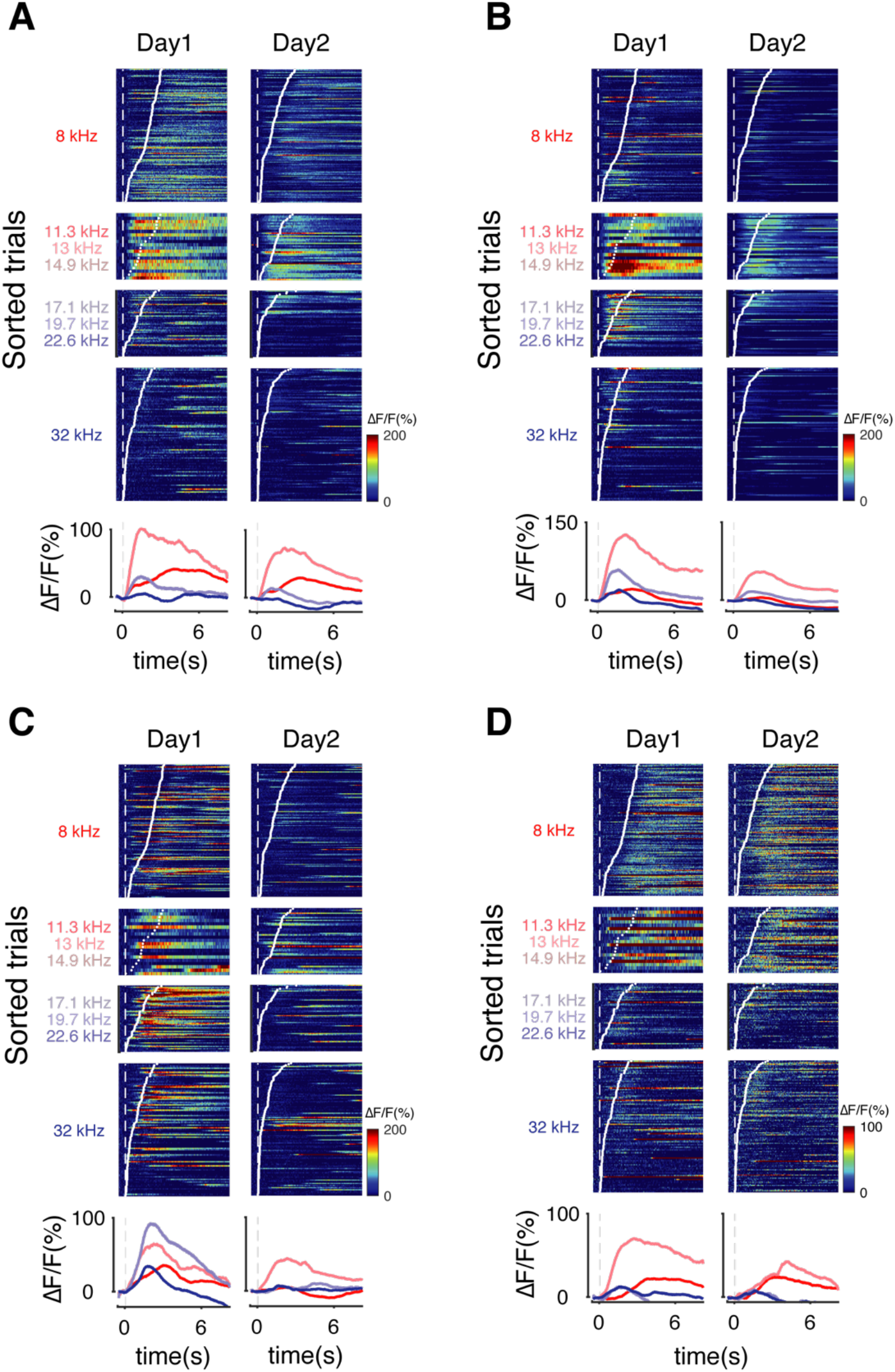
Extra example neurons showing preferred selectivity to test stimuli on day 1 but reduced in the following session.

**Fig. S5.**
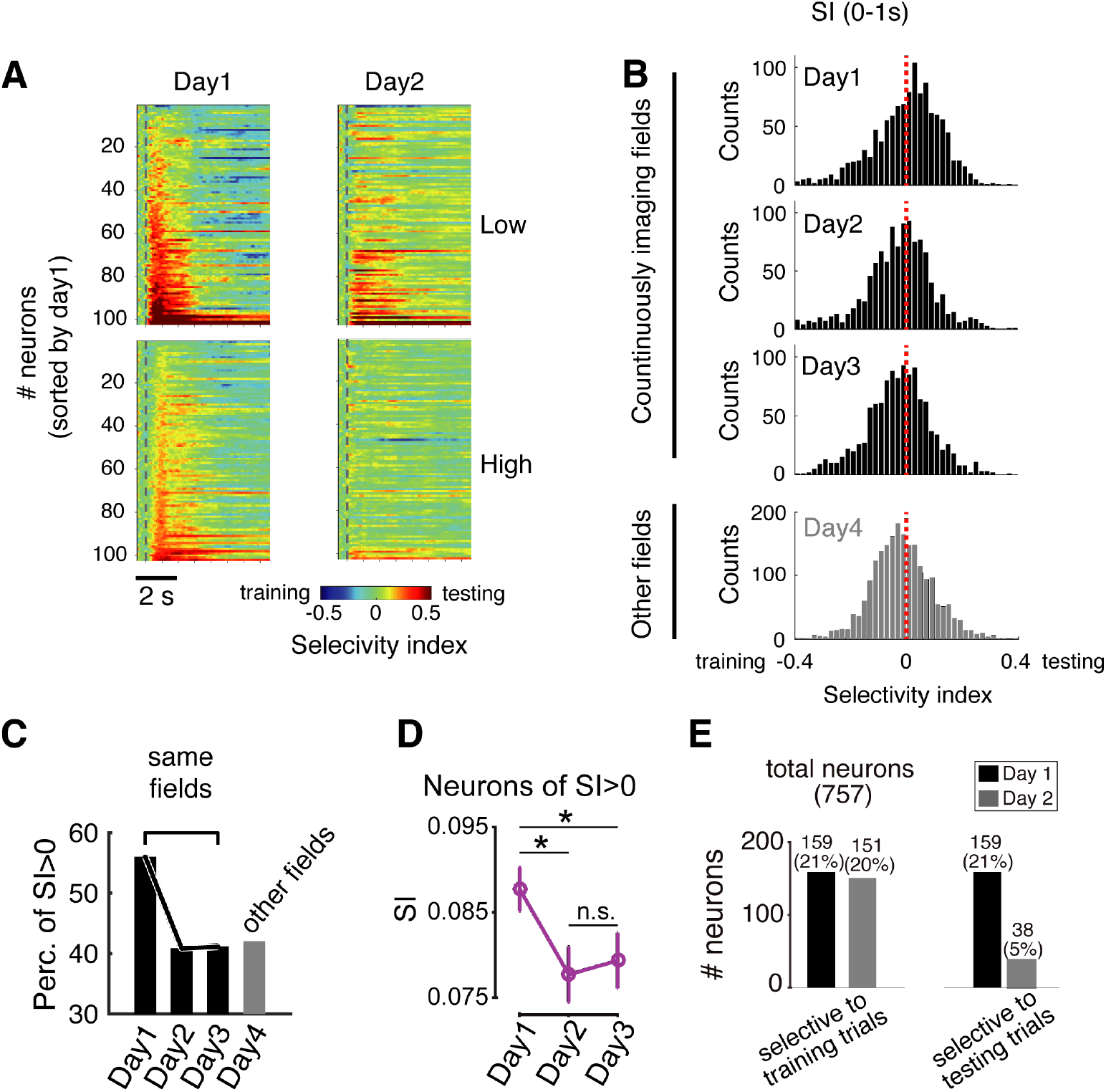
Changes in selectivity for training vs. test stimuli in longitudinally imaged neuron populations. **(A)** Selectivity index for training vs. test stimuli across trial time in simultaneously imaged PPC neurons from an example imaging field across two consecutive days. Upper, data from trials presenting tones of “low” frequency category. Lower, data from trials presenting tones of “high” frequency category. Dash line, stimulus onset. **(B)** Distribution of SI for training vs. testing across four days. Data in day 1 to day 3 were from the same population of neurons, *n* (mice) = 4, *n* (neurons) = 582. Data on day 4 were from neurons in different imaging fields in the same 4 mice, *n* (neurons) = 1055. Only on day 1 was the population selectivity significant biased to testing trials. **(C)** Percentage of neurons showing SI > 0 across 4 days following introduction of test stimuli. **(D)** Selectivity index for test and training stimuli across 3 days for neurons preferring test stimui on day 1. **(E)** Fraction of neurons responded more strongly (*t*-test) training stimuli vs test stimuli on day 1 and day 2.

**Fig. S6.**
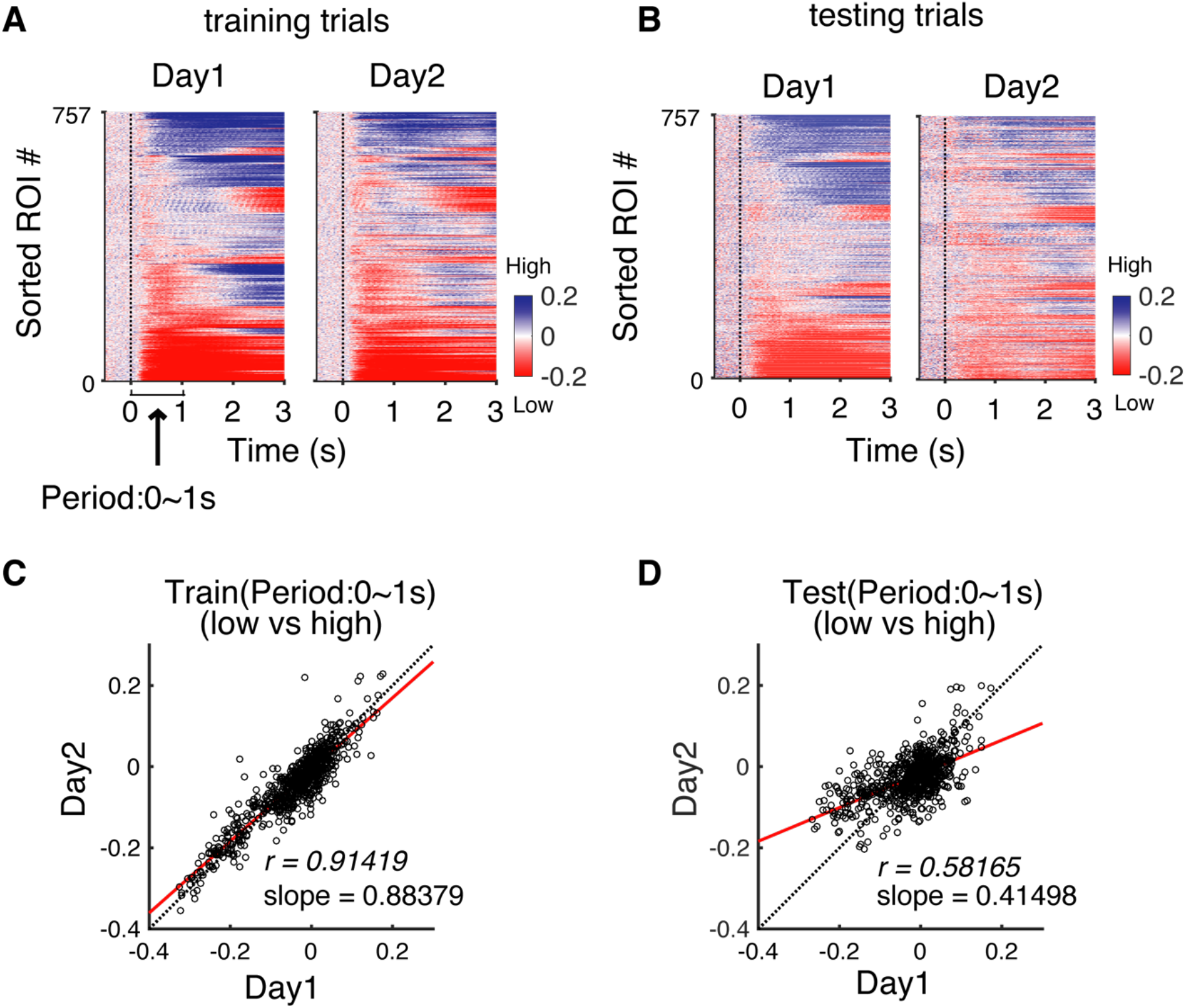
Selectivity for “low” vs. “high” stimulus categories in PPC neuron populations across consecutive days. **(A)** Selectivity index of “low” vs. “high” categories in training trials over trial time in all neurons in two consecutive days. Neurons were sorted by SI values in sequential time windows on day 1. Note that the pattern of SI is preserved in day 2, even using the same neuronal sorting as in day 1. **(B)** Similar in **(A)** for SI computed using testing trials. **(C)** Correlation of SI value in the time window of 1 s after stimulus onset computed using training trials (*P* < 0.0001). **(D)** Similar in **(C)** using data from testing trials (*P* < 0.0001).

**Fig. S7.**
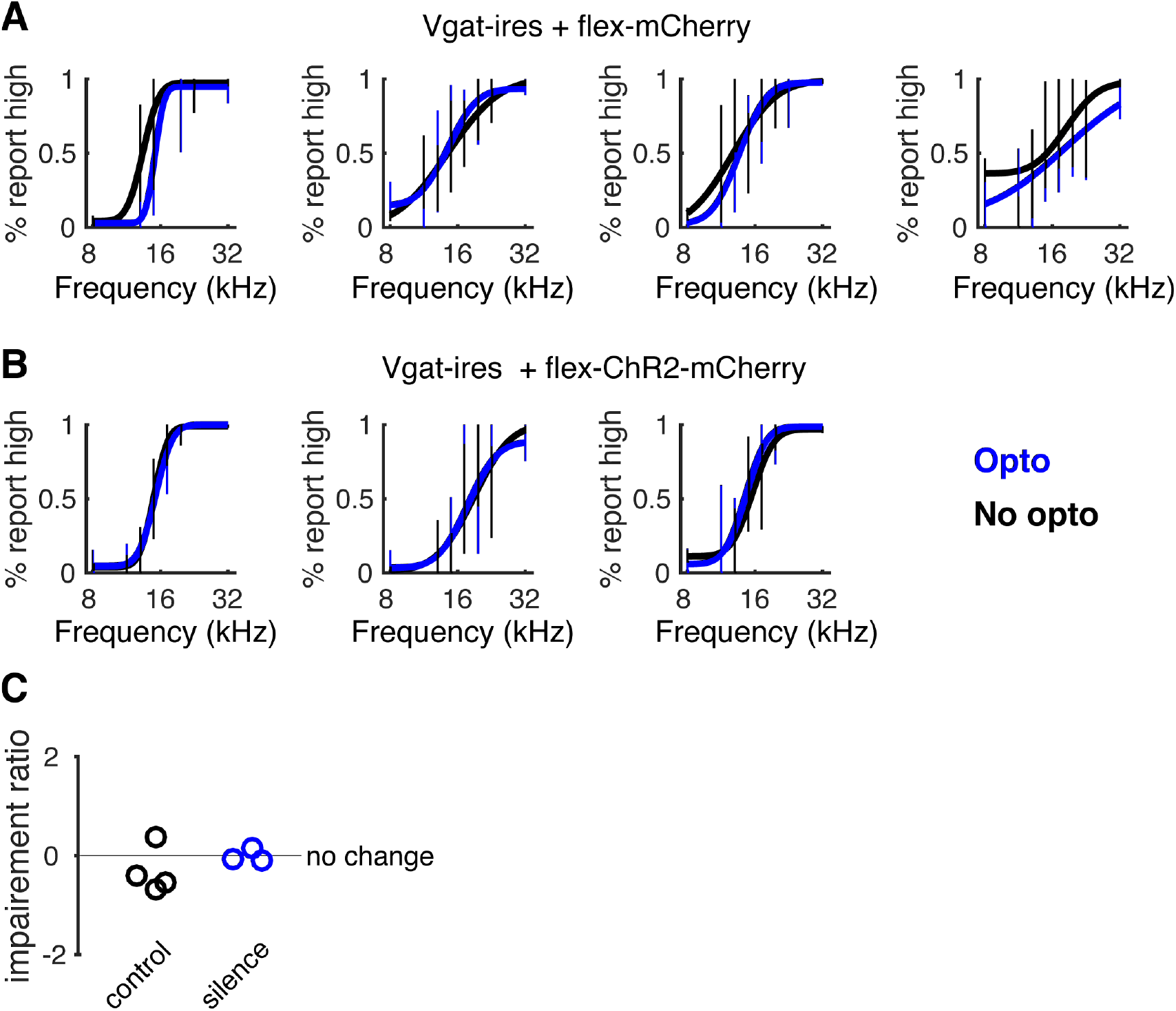
PPC activity in partial of testing trials is sufficient for category learning. **(A)** Psychometric curves for individual animals of the control group, expressed mCherry in PPC. Photostimulation was delivered in 50% of the testing trials, and in 10% of the training trials. Blue curve represents psychometric curves for photo stimulated trials. **(B)** Same plot as in a, for individual animals of the photoinhibition group, with PPC silenced in 50% of the testing trials, and in 10% of the training trials. C. Summary of the mean impairment ratio, (*control-silence) / control*, for data shown in a and (B) Control, *P* = 0.27. Partial silencing, *P* = 0.97, one sample *t*-test.

**Fig. S8.**
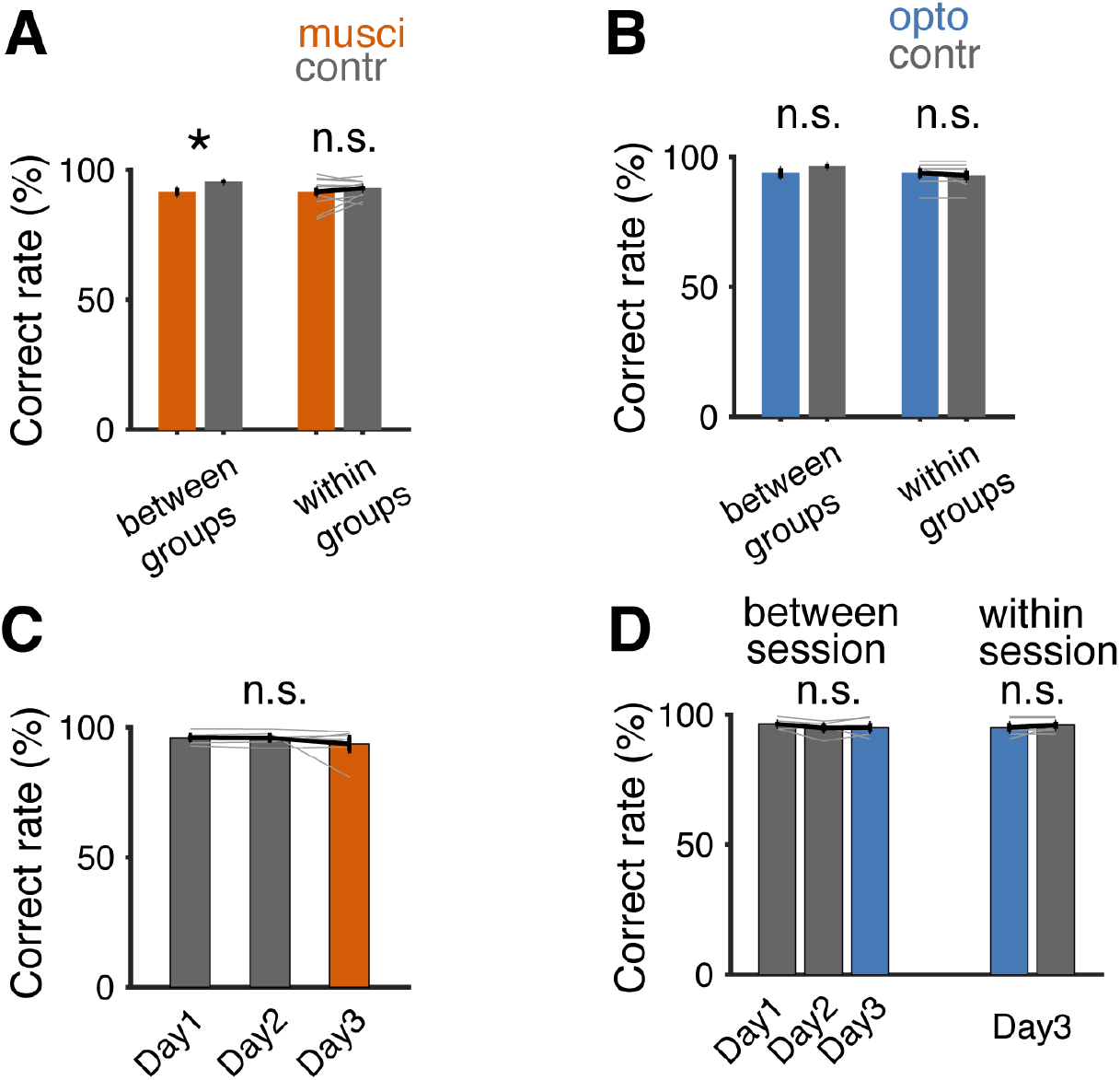
Effects of PPC inactivation on behavioral performance for well-trained tones. **(A)** Behavioral performance for training tones during PPC inactivation by muscimol on the first day introducing test stimuli (*n* = 13), compared with control group (injected with saline, left, *n* = 14, two-sample *t*-test, *P* < 0.05) and compared within group (same animals on the 2^nd^ day when injected with saline, right, *n* = 13, paired *t*-test, *P* > 0.05). Bars, mean ± s.e.m. Light lines, individual mice. **(B)** As in (A) for photoinhibition experiments. Left, comparison between photoinhibition group (blue light + mask, *n* = 7) with control group (mask only, *n* = 6), two-sample *t*-test, *P* > 0.05. Right, comparison for the same animals on day 1 with photoinhibition with day 2 with light mask. *P* > 0.05, paired *t*-test, *n* = 7. **(C)** Behavioral performance for training tones of with muscimol silencing on the 3rd day/session following introduction of test stimuli. *P* > 0.05 between consecutive days, paired *t*-test, *n* = 7. **(D)** Left, similar as in c for photoinhibition, *P* > 0.05 between consecutive days, paired *t*-test, *n* = 5. Right, comparison of behavioral performance for training stimuli between photoinhibition trials (10%) and control trials (90%) on day 3. *P* > 0.05, paired *t*-test, *n* = 5.

**Fig. S9.**
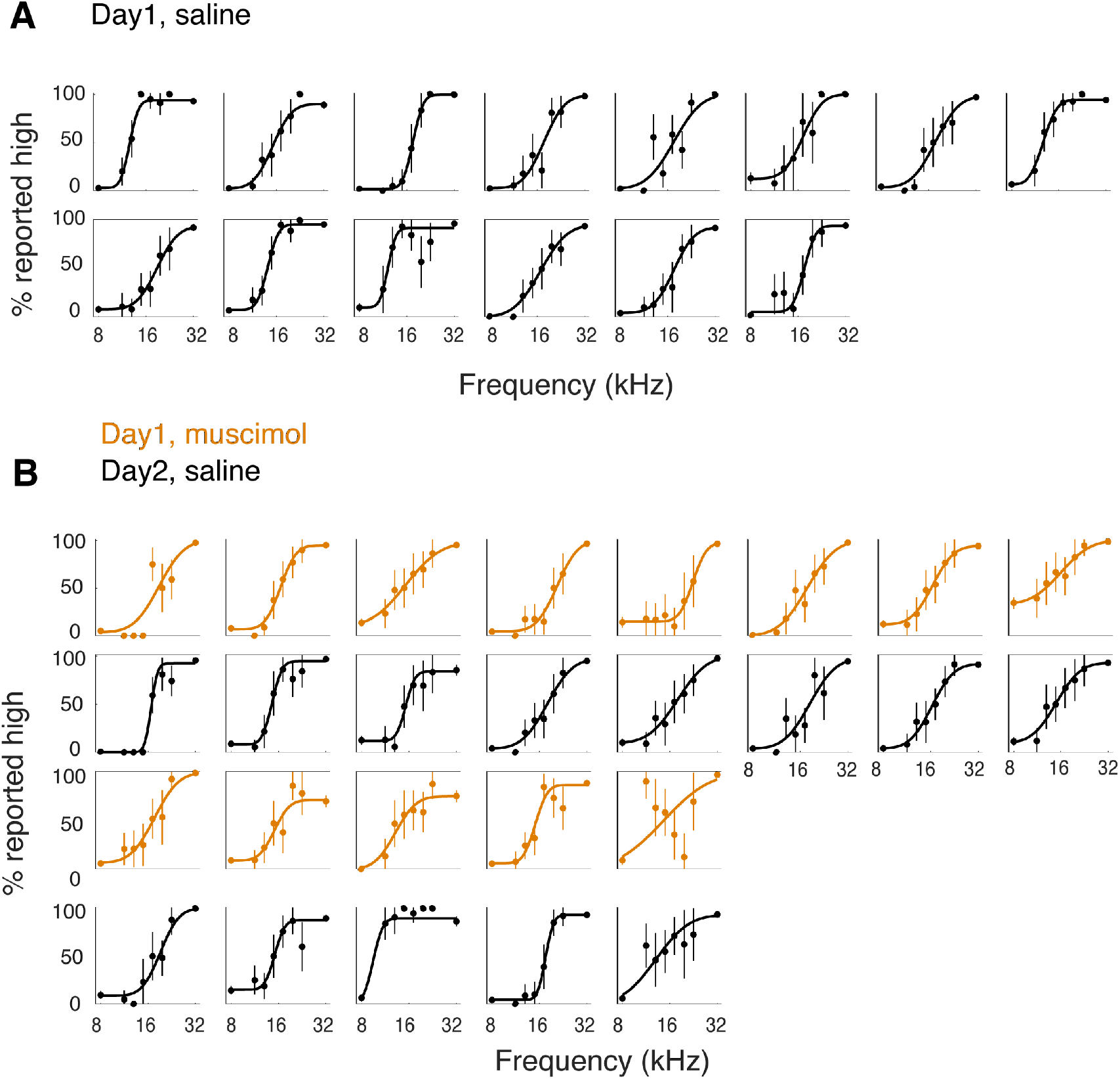
Individual psychometric curves from all animals used for muscimol silencing and saline injection. **(A)** Control group (*n* = 14). Mice received saline injections in PPC on day 1 (first introducing test stimuli). **(B)** PPC silencing group (*n* = 13). Mice received muscimol injections in PPC on day 1, and saline injection on day 2. Error bars, 95% confidence interval.

**Fig. S10.**
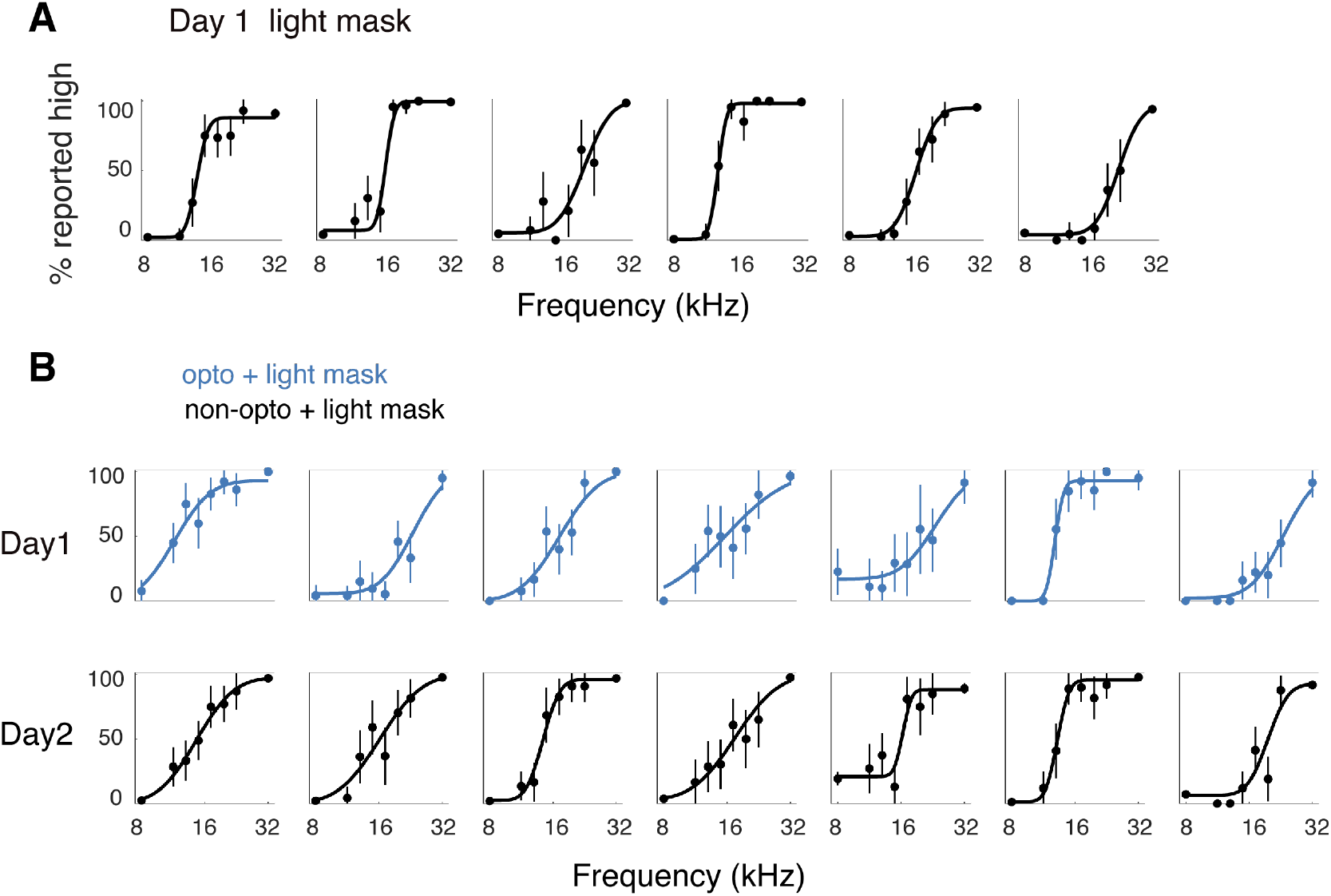
Individual psychometric curves from all animals used for photoinhibition and control. **(A)** Control group (*n* = 6). VGAT-Cre mice received light mask only on day 1 introducing test stimuli. **(B)** Photoinhibition group (*n*= 7). VGAT-Cre mice received blue laser stimulation plus light mask on day 1 and mask only on day 2. Error bars, 95% confidence interval.

**Fig. S11.**
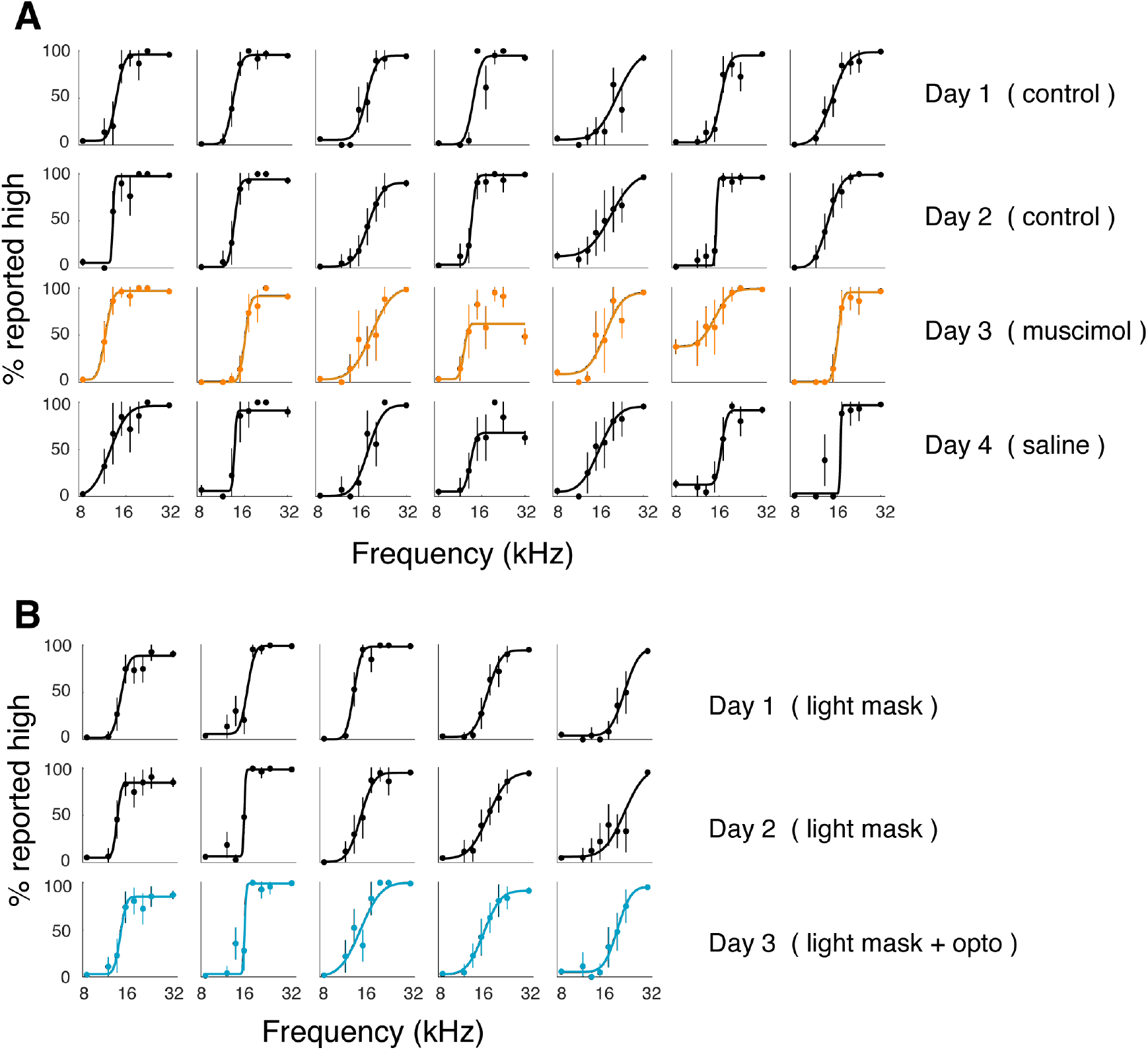
Individual psychometric curves from all animals used for PPC silencing and control on day 3. **(A)** PPC inactivation with muscimol, *n* = 7. **(B)** PPC inactivation with optogenetics, *n* = 5. Error bars, 95% confidence interval.

**Fig. S12.**
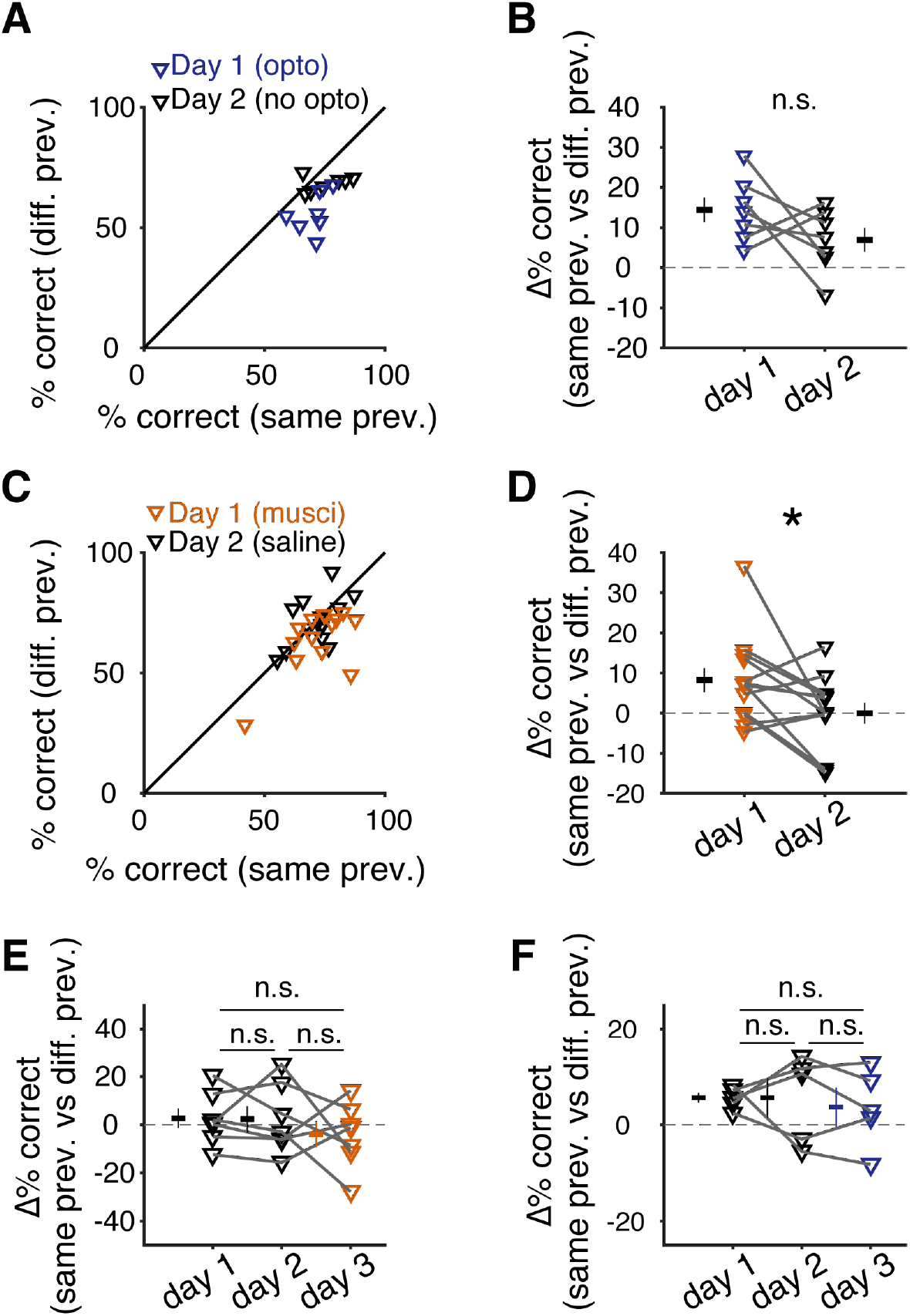
Previous trial history bias during PPC silencing and control conditions in the same animal across different days. (**A** and **B**) Difference in performance for test trials when the previous training trials were of the same or different categories, presented as scatter plot (A) and the values of difference (B) Blue symbols, day 1 with photoinhibition (*P* < 0.05), and day 2 with mask only (*P* > 0.05). *n* = 7, paired *t*-test. (**C** and **D**), Similar as in (A) and (B) for muscimol silencing group (day 1, muscimol, *P* < 0.05; day 2, saline, *P* > 0.05; *n* = 13, paired *t*-test). **(E)** Effect of PPC silencing with muscimol on previous trial history bias on day 3 following introduction of testing trials. Bias is expressed as the difference values as in (B) across 3 days. **(F)** As in **(E)** for the photoinhibition animal group. n.s., not significant.

**Fig. S13.**
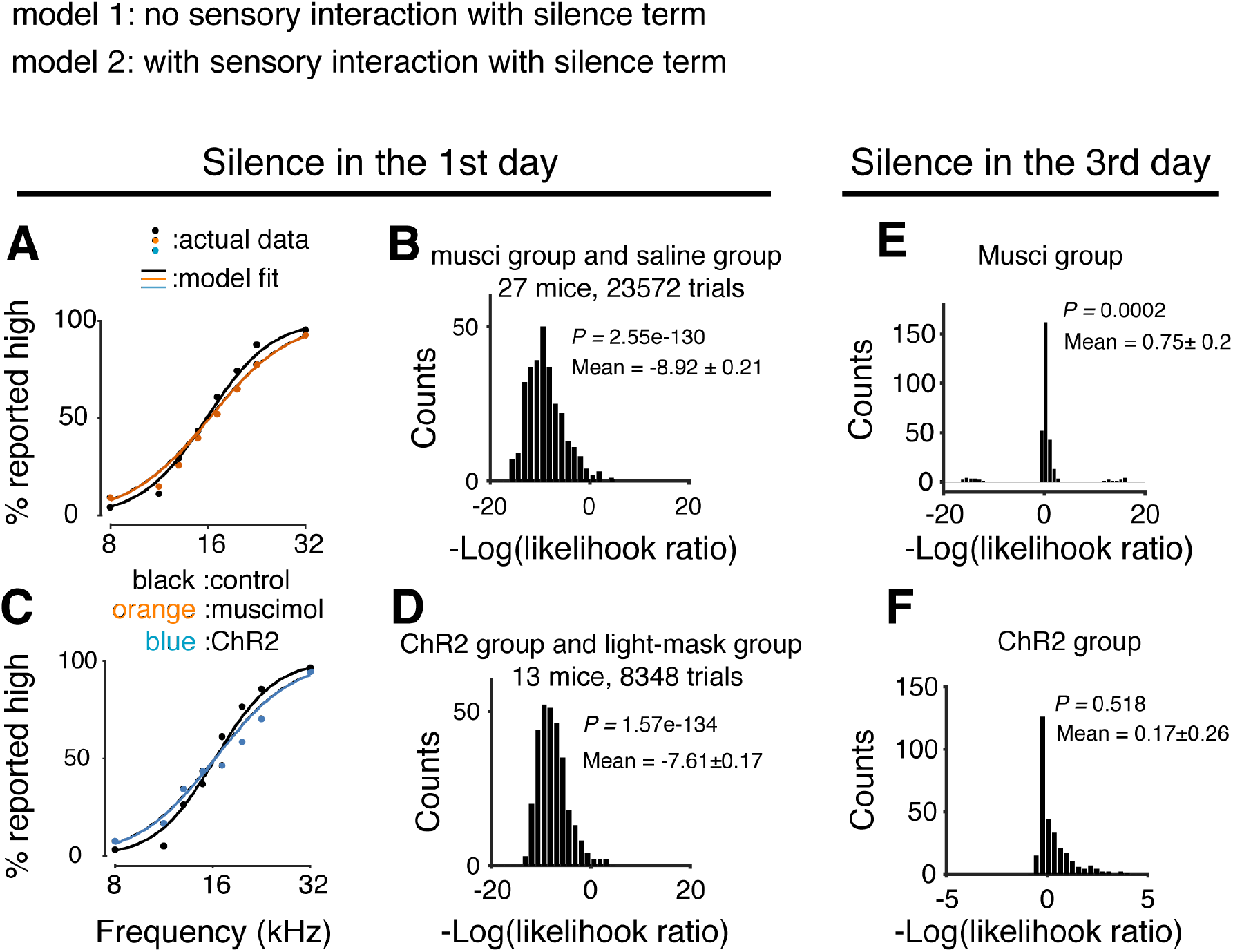
Comparison of GLM models with or without the term of interaction between sensory stimuli and PPC silencing. **(A)** Visualization of slope change in GLM fitting using model with the interaction of stimulus and PPC silencing term. **(B)** Model comparison using likelihood ratio test for models with (model 2) and without (model 1) the interaction of stimulus and silencing term to account for slope change effect during PPC silence on day 1 (first introduction of testing trials). Data from muscimol and saline injection groups (*n* = 27, mean – logLR = −8.92 ± 0.21, *P* < 10^−129^). **(C)** As in (A) for photoinhibition experiments. **(D)** As in (B) for photoinhibition and control groups (*n* = 13, mean –logLR = −7.61 ± 0.17, *P* < 10^−133^). **(E)** Comparison of model 1 and model 2 to account for slope changes for PPC silencing by muscimol on day 3 (testing trials were introduced on day 1). Data from the same animal group on day 1 (control) and day 3 (muscimol injection). *n* = 7, mean –logLR = 0.75 ± 0.2, *P* < 0.001. **(F)** As in (E) for photoinhibition on day 3. *n* = 5, mean –logLR = 0.17 ± 0.26, *P* >0.05. Significance test, paired *t*-test.

**Fig. S14.**
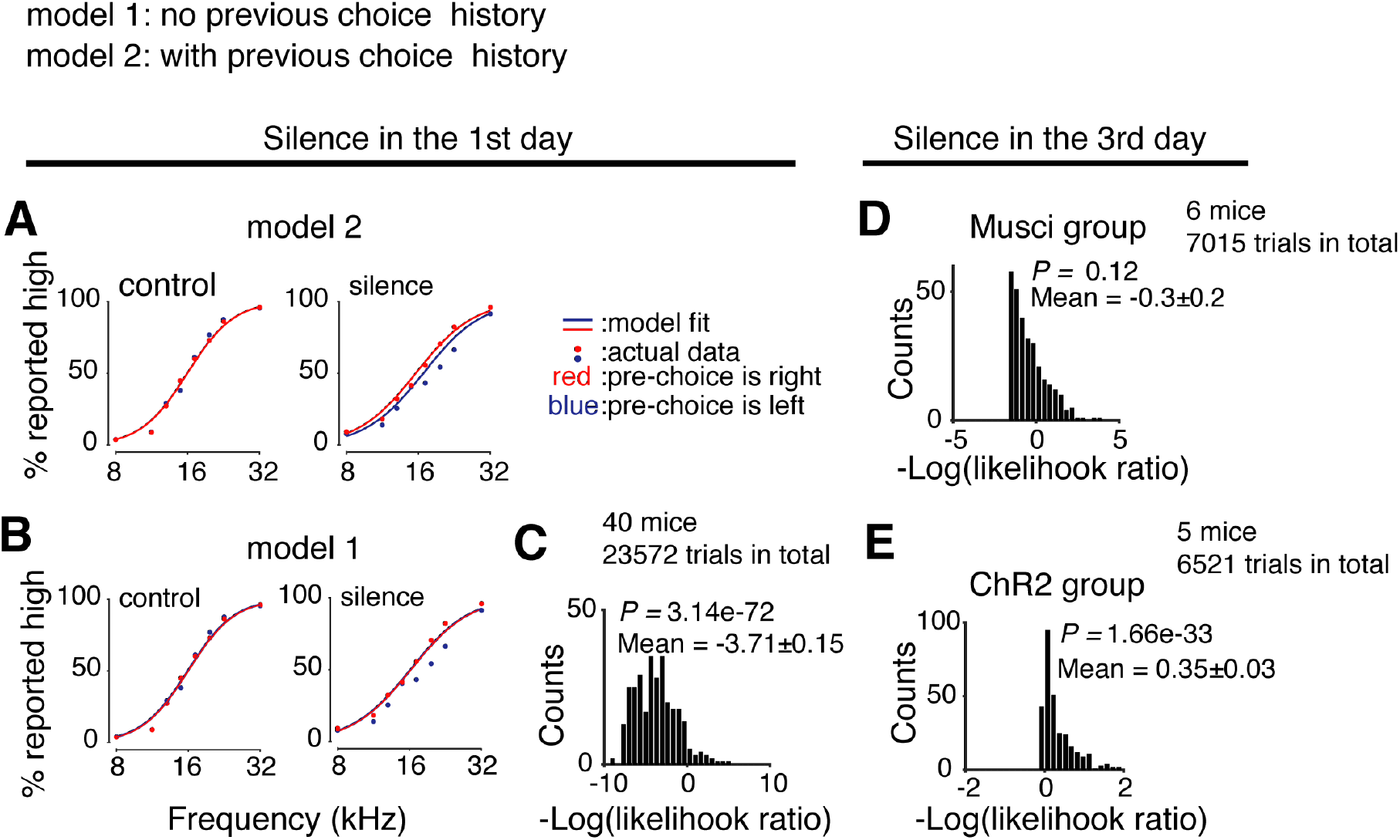
Comparison of GLM models with or without the term of interaction between PPC silencing and previous trial bias. **(A)** Visualization of previous trial choice bias in GLM fitting using model 1 (without the interaction of previous trial choice and PPC silence term). Red, data from trials with previous choice being right licking. Blue, data from trials with previous choice being left licking. Left, data from control condition. Right, data from PPC silencing (including both muscimol and photoinhibition). **(B)** As in (A), using model 2 (with the interaction of previous trial choice and PPC silencing term). Note a clear shifting in the data points during PPC silencing that is better captured by model 2. **(C)** Model comparison using likelihood ratio test for models 1 and 2 in accounting for previous trial choice bias effect when PPC was silenced on day 1 introducing testing trials. Data from all groups (*n* = 40, mean –logLR = −3.71 ± 0.15, P < 10^−71^, paired *t*-test). (**D** and **E**) Model comparison using likelihood ratio test for models 1 and 2 fitted to the data of PPC silencing on day 3, using muscimol (D) mean –logLR = −0.3 ± 0.2, *P* = 0.12, *n* = 6, and photoinhibition **(E)** mean –logLR = 0.35 ± 0.03, *P* < 10^−32^, *n* = 5. Paired *t*-test.

**Fig. S15.**
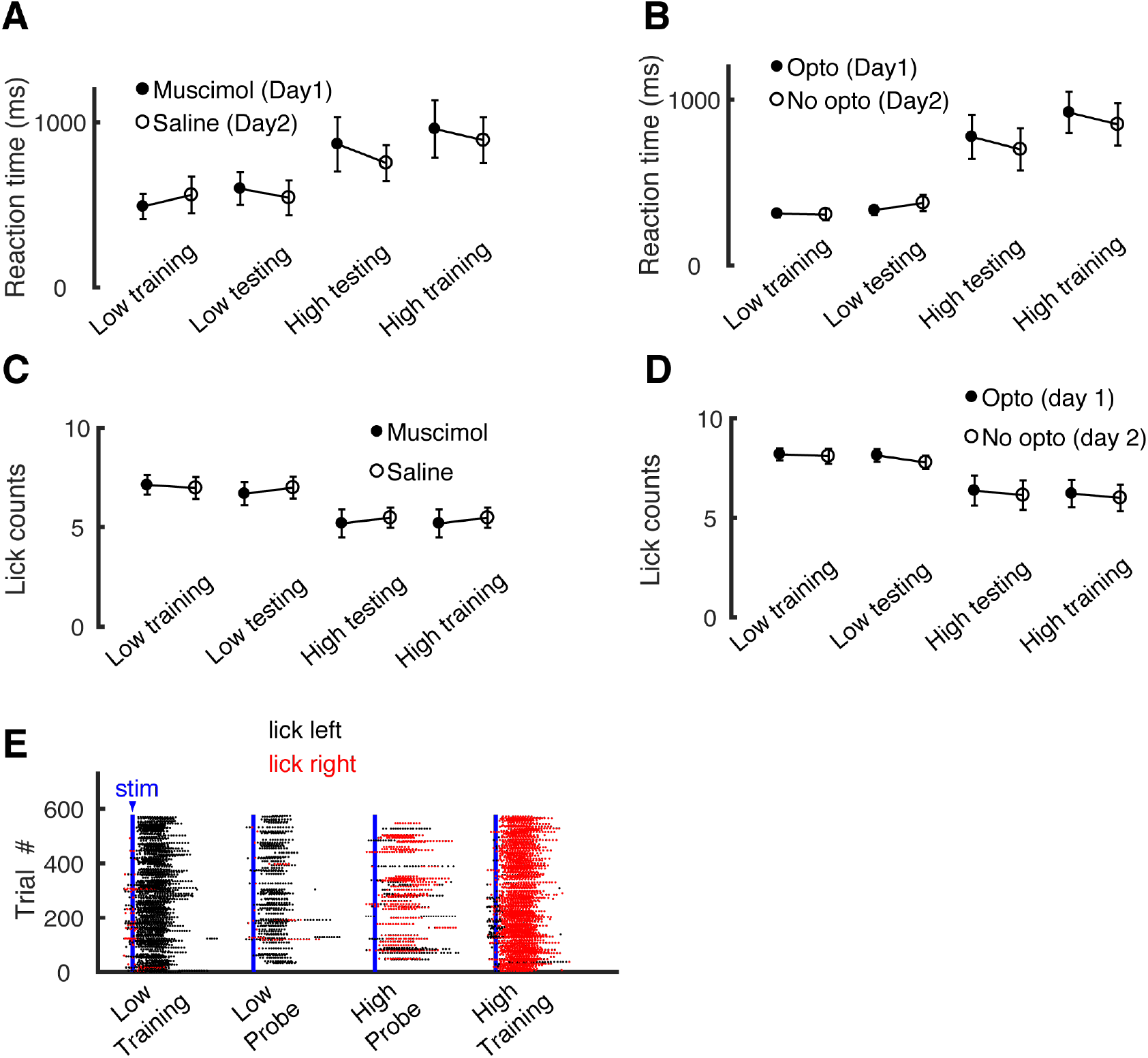
Effect of PPC silencing on reaction time and licking actions. **(A)** Comparison of reaction time (first lick after stimulus onset) between muscimol injection (day 1) and saline injection (day 2) conditions of the same animals. Data were segregated based on trial types. *P* > 0.05 for all comparisons, *n* = 13. **(B)** Similar in (A) for data in photoinhibition of PPC. *P* > 0.05, *n* = 7. **(C)** Similar plot as in (A) for mean lick number per trial in 1.5 s window after stimulus onset, compared between muscimol silencing and control conditions. *P* > 0.05 for all comparisons, *n* = 13. **(D)** Similar in (C) compared between photoinhibition and control conditions. *P* > 0.05 for all comparisons, *n* = 7. Paired *t*-test. **(E)** Raster plot of lick times in an example session with PPC silenced.

**Table S1.** Overlap of category selectivity. Summary of fraction of neurons showing response selectivity to different behavioral variables. Neurons are classified based on alignment of response time regarding to stimulus onset time, answer lick time, occurrence in correct trials and error trials respectively, or occurrence in subset of trials with certain tone frequencies. Neurons with multiple response peaks are classified as mixed selectivity, e.g., “Sensory + choice”. Some neurons show large responses late in trials but still correlate with stimulus or choice are classified as “Sensory + late response” or “Choice + late response”.

|  | n / Total | Percentage |
| --- | --- | --- |
| No selectivity | 29/758 | 4% |
| Sensory | 21/758 | 3% |
| Choice | 174/758 | 23% |
| Late response | 161/758 | 21% |
| Sensory + choice | 156/758 | 21% |
| Sensory + late response | 93/758 | 12% |
| Choice + late response | 268/758 | 35% |
| Sensory + choice+ late response | 72/758 | 9% |

